# Smooth Muscle Cell Death Drives an Osteochondrogenic Phenotype and Severe Proximal Vascular Disease in Progeria

**DOI:** 10.1101/2023.01.10.523266

**Authors:** Sae-Il Murtada, Yuki Kawamura, Cristina Cavinato, Mo Wang, Abhay B. Ramachandra, Bart Spronck, George Tellides, Jay D. Humphrey

## Abstract

Hutchinson-Gilford Progeria Syndrome results in rapid aging and severe cardiovascular sequelae that accelerate near end of life. We associate progressive deterioration of arterial structure and function with single cell transcriptional changes, which reveals a rapid disease process in proximal elastic arteries that largely spares distal muscular arteries. These data suggest a novel sequence of progressive vascular disease in progeria: initial extracellular matrix remodeling followed by mechanical stress-induced smooth muscle cell death in proximal arteries, leading a subset of remnant smooth muscle cells to an osteochondrogenic phenotypic modulation that results in an accumulation of proteoglycans that thickens the wall and increases pulse wave velocity, with late calcification exacerbating these effects. Increased pulse wave velocity drives left ventricular diastolic dysfunction, the primary diagnosis in progeria children. Mitigating smooth muscle cell loss / phenotypic modulation promises to have important cardiovascular implications in progeria patients.

## INTRODUCTION

The systemic arterial tree normally develops to have a differential structure and function along its length that optimizes hemodynamics and cardiovascular physiology, with large (elastic) arteries conferring appropriate compliance and resilience and medium-sized (muscular) arteries enabling high-level vasoactive control. While exhibiting important differential biomechanical properties, these vessels also experience differential disease manifestations, as in hypertension and aging wherein elastic arteries stiffen more than muscular arteries (Laurent & Boutouyrie, 2015). Hutchinson-Gilford Progeria Syndrome (HGPS, or simply progeria) is an ultra-rare condition resulting from a mutation to the gene (*LMNA*) that encodes the nuclear scaffolding protein lamin-A (De Sandre-Giovannoli et al., 2003; Eriksson et al., 2003); it leads to diverse clinical sequelae and premature death. Early cardiovascular studies of progeria patients suggested a key role of atherosclerosis in muscular (coronary) arteries (Olive et al., 2010) whereas a recent clinical study found left ventricular diastolic dysfunction to be the most common diagnosis (Prakash et al., 2018), consistent with a prior report that implicated a stiffening of the aorta (elastic artery) that was reflected by an increased pulse wave velocity (Gerhard-Herman et al., 2012). Importantly, this progeria phenotype progresses rapidly near end of life.

Notwithstanding a growing clinical data base (Gordon et al., 2012, 2014, 2018), mouse models of progeria have emerged as critical to increasing understanding (Capell et al., 2008; Osorio et al., 2011). Nevertheless, information on the expected progressive deterioration of arterial structure and function remains limited. Here, we biomechanically phenotype five representative systemic arteries – the ascending (ATA) and descending (DTA) thoracic aorta, the common (CCA) and internal (ICA) carotid artery, and the second branch of the mesenteric artery (MA) – and transcriptionally profile the aorta from a well-accepted mouse model of progeria (*Lmna^G609G/G609G^*) at four critical times of vascular development / disease progression: postnatal days P42, P100, P140, and P168. The ATA and DTA are key elastic energy storing arteries that augment blood flow via a Windkessel effect; the CCA is somewhat of a transitional artery, though more elastic; the ICA and MA are muscular arteries. Importantly, cell damage is highly mechano-dependent in progeria (Broers et al., 2004; Verstraeten et al., 2008) and intramural mechanical stress differs markedly along the normal arterial tree, with values in the adult wild-type (WT) mouse greater than 250 kPa in the ATA and DTA and less than 100 kPa in the ICA and MA, with that in the CCA between. Material stiffness increases linearly with intramural stress in the murine aorta (Humphrey and Tellides, 2019). Normal levels of lamin-A expression increase with increasing stiffness of the matrix in which the cells reside (Swift et al., 2013), presumably to increasingly stress-shield nuclear contents and prevent mechanically induced DNA damage. Lamin-A is highly expressed in the highly stressed aorta (Kim et al., 2018). We hypothesized that progeria should lead to differential progression of arterial disease due, in part, to normal regional variations in intramural stress and stiffness from elastic to muscular arteries. Because of a relatively early accumulation of mural proteoglycans as well as late-stage emergence of medial calcification, particularly in the aorta, we used bulk and single-cell RNA sequencing to examine cell phenotypic changes, particularly in the smooth muscle cells (SMCs) given the dramatic histological changes observed in the medial layer.

## RESULTS

### Progressive microstructural abnormalities emerge in elastic arteries in progeria

Focusing first on the period near end of life in these *Lmna^G609G/G609G^* progeria mice (P168), standard histological examinations revealed regional differences in accumulated proteoglycans, which tend to thicken the wall, plus late mural calcification, especially in the DTA (Fig. 1A). Focusing further on the descending aorta, histology revealed increases in and straightening of mural collagens (Fig. 1B), especially in the adventitia, which was also seen under *in vivo* relevant conditions using multiphoton microscopy (Fig. 1C). These changes in adventitial collagen are inconsistent with homeostatic turnover. Multiphoton microscopy also uncovered a significant (~67%) loss of medial (SMCs) cells and a trend toward a loss of intimal (endothelial) cells in the DTA in progeria at P168 (Fig. 2A,B). This SMC drop-out was confirmed in the DTA at P168 by TUNEL immuno-staining for apoptotic cells (Fig. 2C,D), noting that the SMC nuclei appeared distorted and irregular in shape (Supplemental Fig. S1).

**Fig. 1.**
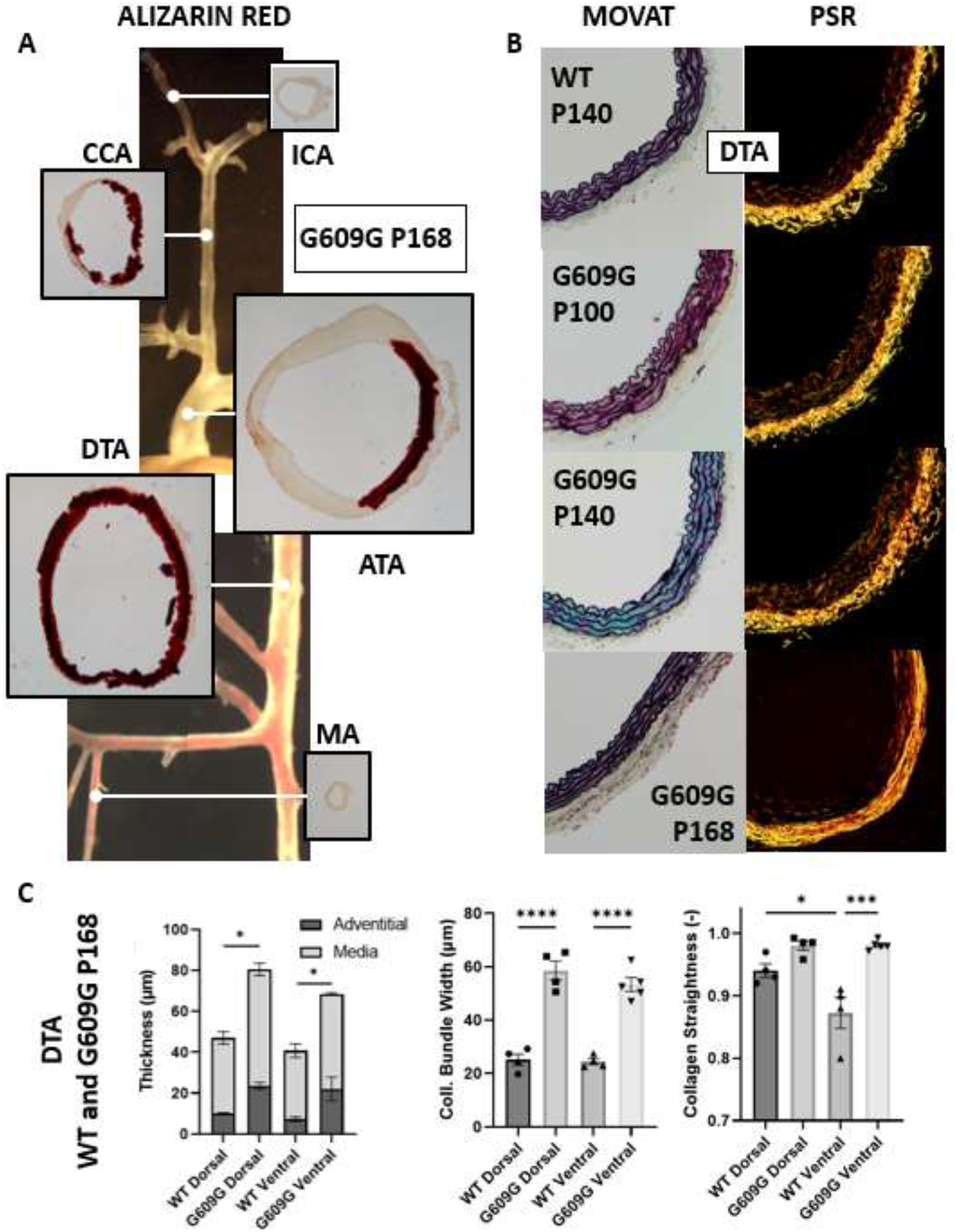
(A) Background: Gross stitched photographs of the five arterial segments of interest (ascending thoracic aorta - ATA, descending thoracic aorta - DTA, common carotid artery - CCA, internal carotid artery - ICA, and mesenteric artery - MA) in the progeria mouse (G609G) at postnatal day P168; Foreground: associated illustrative histological cross-sections (Alizarin red) showing mural calcification at P168, if present. (B) Illustrative histological cross-sections (bright-field Movat pentachrome - MOVAT, dark-field picrosirius red - PSR) of the DTA comparing wild-type (WT) at P140 vs. progressive changes in progeria from postnatal day P100 to P140 to P168. Note, in particular, the marked accumulation of GAGs at P140 and the increased thickening of the adventitia at P168 with associated straightening of the collagen fibers in progeria. (C) Further quantitation of histology for the DTA at P168 using multiphoton microscopy: differences in wall thickness by layer (media and adventitia) as well as, via second harmonic generation, differences in adventitial collagen fiber bundle width and straightness.

**Fig. 2.**
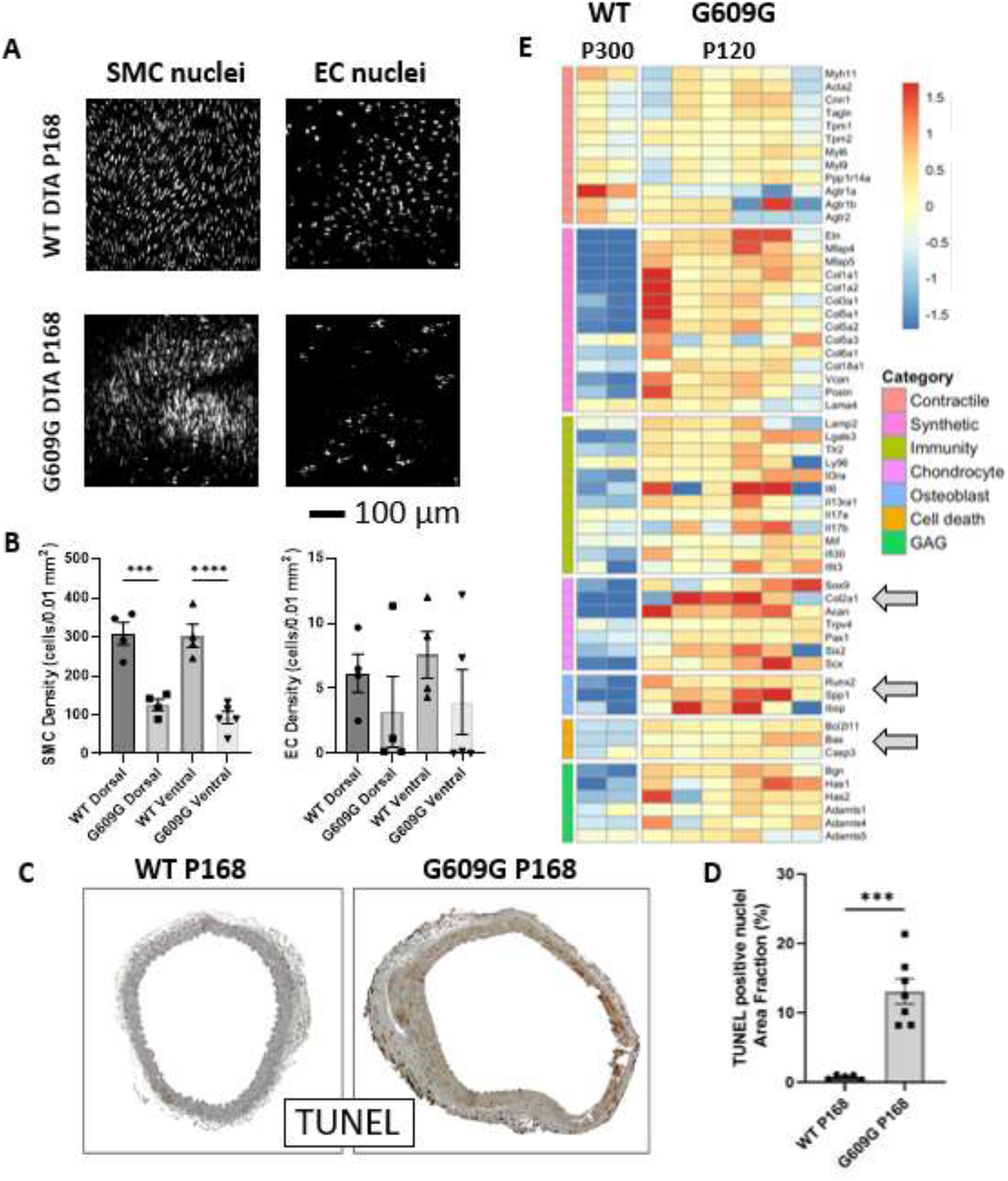
(A) Representative results from multiphoton microscopy comparing the density of cell nuclei both within the media and along the luminal surface of the descending thoracic aorta (DTA) at P168 for WT and G609G. See also Supplemental Fig. S1, which shows better the distorted shapes of the smooth muscle cell nuclei in progeria. (B) Quantitative summary of nuclei density for all images similar to panel A, except with data shown for both the dorsal (upper side, or toward the back) and ventral aspects. (C) TUNEL immunostaining (brown) revealing a significant increase in apoptosis in the DTA at P168 in progeria relative to age-matched WT control, (D) with quantification. (E) Bulk RNA-seq comparing transcripts for wild-type (WT at an aged postnatal day P300, *n*=2) and progeria (G609G at P120, *n*=6) aortas, noting in particular the marked increase in synthetic, degradative (immunity), chondrogenic, osteogenic, and apoptotic markers in progeria. The arrows not, in particular, the increase in apoptotic transcripts (e.g., Bax) as well as chondrogenic (Sox9, Acan, Col2a1) and osteogenic (Runx2, Spp1) transcripts.

### RNA sequencing reveals a progressive phenotypic modulation of aortic SMCs in progeria

To understand better the reasons for distinctive changes in aortic composition in progeria, we first performed bulk RNA-sequencing on the abdominal aorta, which exhibits a phenotype similar to the thoracic aorta in progeria (Murtada et al., 2020). Contrasting results for naturally aged WT and progeria aortas well prior to end-of-life (P300 and P120, respectively) revealed dramatic differences in many transcripts (Fig. 2E). Although markers for a contractile SMC phenotype (e.g., Myh11, Tgln) tended not to differ dramatically at this age, there was a marked increase in transcripts for extracellular matrix synthesis (e.g., Col1a1, Col3a1, Vcan) as well as those for apoptosis (Bcl2l11, Bax, Casp3) and an osteochondrogenic phenotype (e.g., Sox9, Col2a1, Acan, Runx2, Spp1) – see arrows in panel E of Fig. 2. We thus performed single cell RNA sequencing (scRNA-seq) on whole (thoracic + abdominal) aortas from progeria mice at postnatal days P100, P140, and P168 to compare with adult WT. Twenty clusters emerged (Fig. S2), including those for endothelial cells, SMCs, fibroblasts, and inflammatory cells, including macrophages. Notwithstanding the tremendous information available, we again focused on SMCs given the biomechanical significance of the media in elastic arteries and dramatic histological changes in the media in these vessels in progeria. Importantly, SMC clusters not only included cells that expressed contractile (e.g., Acta2, Tgln) and synthetic (e.g., Col1a1, Col3a1) phenotypic markers, they also expressed chondrocytic markers (Sox9, Acan) – Figs. 3A-H, S3A-C. Markers for an osteogenic phenotype (e.g., Runx2, Spp1) were expressed in one SMC cluster while also associating with a macrophage cluster. When contrasting these changes in SMCs at different ages (Fig. S3B,C), there appeared to be a slight spike in markers for a contractile phenotype at P140 prior to a spike in synthetic phenotype at P168 in progeria. Importantly, a few cells expressed markers of an osteochondrogenic phenotype in the P140 WT or P100 progeria aortas, but there was a progressive rise in progeria, peaking at P140 (Figs. 3I, S4A,B). A gene ontology analysis of the scRNA-seq results for SMCs revealed an abnormal bone formation process (Fig. S4C), consistent with histological evidence of marked calcification of the DTA at P168 (Fig. S4D,E).

**Fig. 3.**
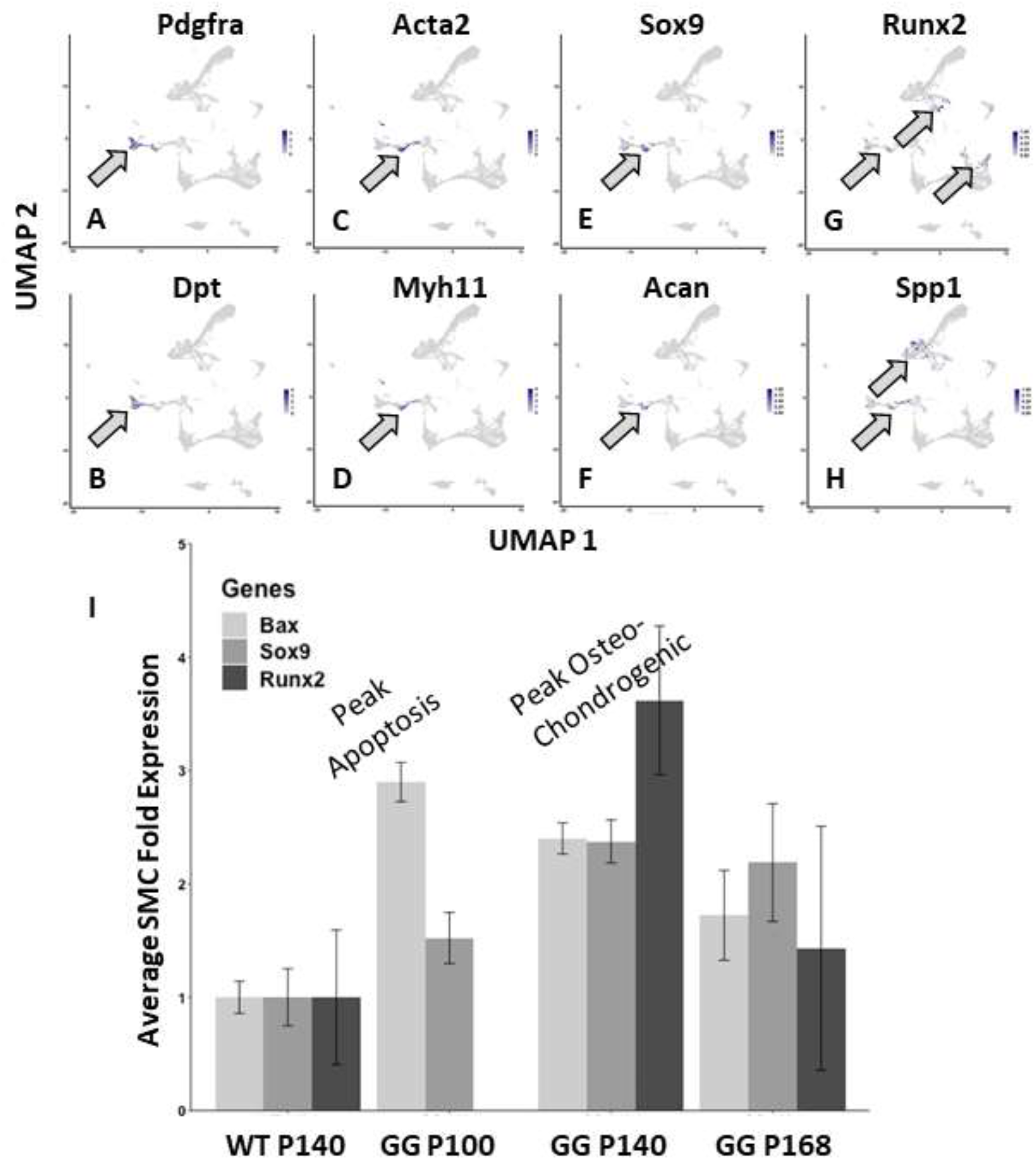
Single-cell RNAseq for the progeria aorta for postnatal days P100-P168. (A,B) Markers for adventitial fibroblasts, (C,D) markers for vascular smooth muscle cells, (E,F) chondrogenic markers, and (G,H) osteogenic markers. Note that the chondrogenic and osteogenic markers co-localize with the smooth muscle markers, though osteogenic markers also co-localize with inflammatory cell markers (see Fig. S2). (I) Time-course of changes in three markers of interest: Bax (gene for Bcl-2 Associated X-protein, which is pro-apoptotic), Sox9 (SRY-Box-Transcription factor 9, which is chondrogenic), Runx2 (Runx Family Transcription Factor 2, which is osteogenic – also known as Cbfa1, core binding factor subunit alpha). Note that apoptotic markers peak early, at P100, whereas the osteo-condrogenic markers peak later, at P140. See also Supplemental Fig. S3 and S4.

### The biomechanical phenotype deteriorates more in elastic than in muscular arteries in progeria

Changes in mural composition are expected to manifest at the tissue level as changes in geometric metrics and material properties. Age-dependent changes in inner radius and wall thickness impact multiple key biomechanical metrics, including wall stress and pulse wave velocity (PWV), a measure of structural stiffness of the wall. In contrast to changes seen during normal arterial development (Murtada et al., 2021a), inner radius decreased from P42 to P168 in progeria in the ATA, DTA, and CCA while not changing appreciably in the ICA or MA (Fig. S5A-E). Wall thickness also exhibited an unnatural trend, increasing appreciably in all five segments in progeria (Fig. S5F-J). The ratio of inner radius *a* to wall thickness *h* is fundamental to calculating mean circumferential intramural stress (*σ*_*θ*_ = *Pa*/*h*, where *P* is pressure) and pulse wave velocity (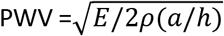, where *E* is the circumferential material stiffness and *ρ* the mass density of the blood). This important geometric ratio, *a*/*h*, decreased with age in all five segments, but especially in the ATA, DTA, and CCA (Fig. S5K-O).

Passive pressure-diameter data collected at specimen-specific values of axial stretch (Fig. S6A-E) revealed a progressive loss of distensibility (left-ward shift) in the aorta (ATA, DTA) and CCA with increasing age from P42 to P168 in progeria, with more complex changes in the ICA and early but sustained changes in the MA. This perceived loss of distensibility was reflected by progressively lower values of *in vivo* relevant circumferential stretches (Fig. S7A-E). Circumferential material stiffness increases modestly from P42 to steady state values ~P100 in WT arteries (Murtada et al., 2021a), yet complex regionally dependent trends emerged in progeria. This stiffness tended to remain the same or decrease from P42 to P168 except in the DTA and ICA, which showed late increases; values tended nonetheless to be less than normal (Fig. S8A-E). All arteries are subjected to biaxial loads *in vivo*, and changes in the axial direction are often the first to manifest in adaptations (Humphrey et al., 2009). Findings were generally similar in the axial direction, namely for axial force-extension (structural) behaviors as well as *in vivo* values of axial stretch and axial material stiffness in all five segments at all four ages in progeria. Specifically, there was a progressive decrease in extensibility in progeria as evidenced by the left-ward shift in the axial force-stretch data (Fig. S6F-J). The *in vivo* value of axial stretch tends to reflect the elasticity of the vessel; it decreased markedly in progeria with age, especially in the ATA and CCA (Fig. S7F-J). Axial material stiffness decreased with age in the aorta (ATA, DTA), increased markedly in the CCA and modestly in the ICA, but remained similar in the MA (Fig. S8F-J).

Two of the most important biomechanical metrics in central arteries are the ability to store elastic energy during systole, to be used during diastole to augment blood flow, and PWV, which reflects the impact of the structural stiffness on the speed of propagation of the pulse pressure wave, a key determinant of cardiovascular health and disease (Boutouyrie et al., 2021). Elastic energy storage capacity was lower than normal in all five regions at all four times in progeria (Fig. 4A), but especially in the proximal arteries indicative of an extreme loss of mechanical function. By contrast, PWV increased from P42 to P168 in all segments in progeria (Fig. 4B), but especially in the DTA near end of life (~P168). Numerical values for these and other biomechanical metrics are in Supplemental Tables S1-S5 for P42-P168, with pressure-dependent values provided at systolic and mean arterial pressures and WT values at P168 given as controls.

**Fig. 4.**
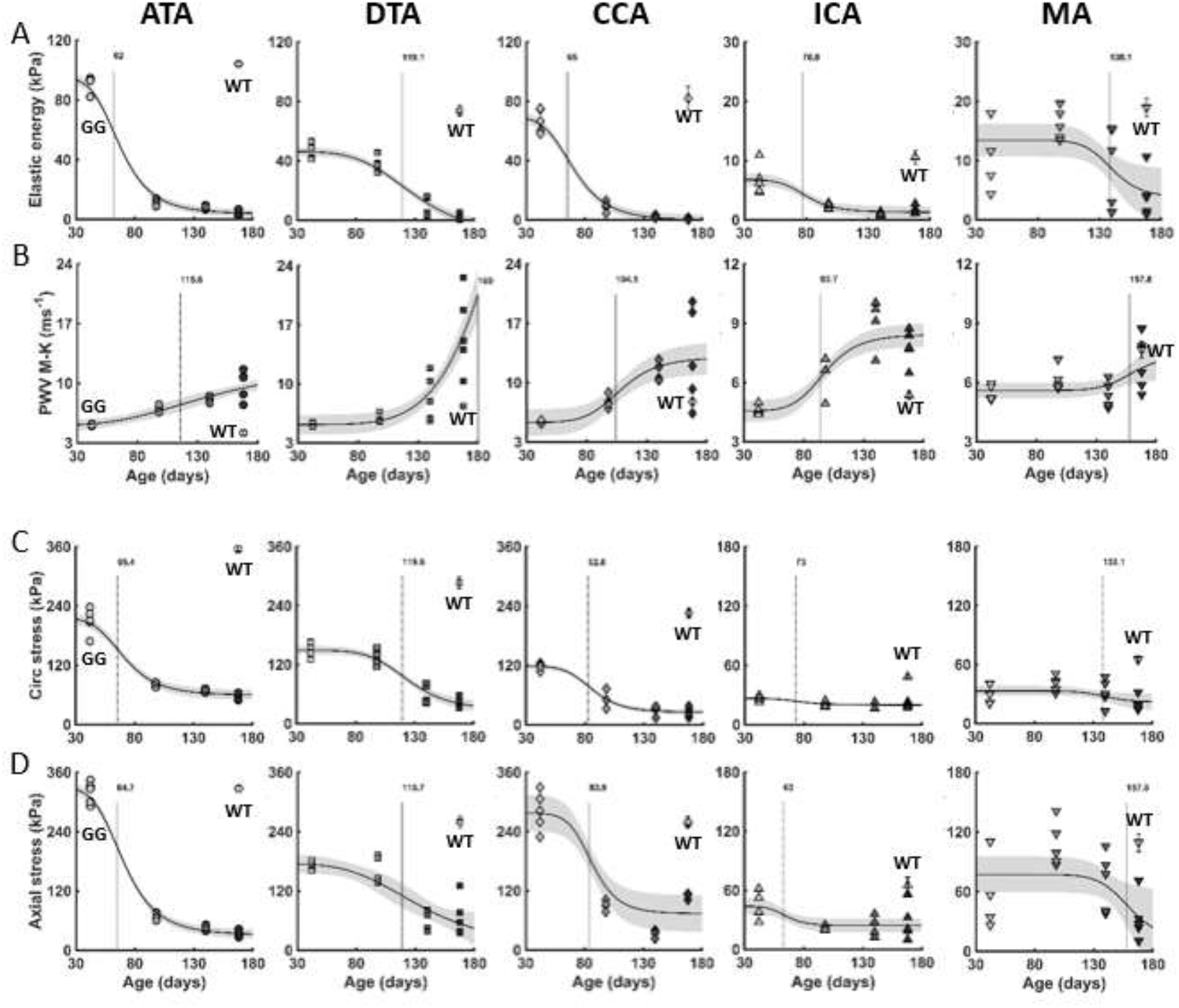
Time-course (from postnatal day P42 to P100, P140 and P168) of key biomechanical metrics in G609G/G609G (GG) progeria mice across five systemic arteries: the ascending (ATA) and descending (DTA) thoracic aorta, the common (CCA) and internal (ICA) carotid artery, and the 2^nd^ branch mesenteric artery (MA). Functional metrics: (A) elastic energy storage capability and (B) local pulse wave velocity (PWV) calculated using the Moens-Korteweg (M-K) equation. Intramural stresses: (C) mean circumferential and (D) mean axial. The open symbol (mean ± SEM) at P168 shows mature wild-type (WT) values for comparison. The vertical dotted line indicates the day at which the rate of change is maximal. *n* = 4-5 per region per time. See also Supplemental Tables S1-S6 for quantitative information and Supplemental Fig. S5-S8.

### Biomechanical dysfunction emerges first in elastic arteries in progeria

Given an analytical description of the individual time-courses of change in the geometric and mechanical metrics (Table S6), we computed the age at which each metric changed most rapidly (vertical lines in Figs. 4, S5-S8). Plotting the age at which each metric reached 90% of its near end of life value (at P168) versus the day of maximum rate of change of that variable highlighted the differential regional variations and when they emerged (Fig. S9A-C). As it can be seen, the day at which a maximal rate of change occurred was seen first for circumferential stiffness of the ATA and CCA, both around P50, followed by changes in their inner radius at ~P70 days and wall thickness at ~96 days. The maximum rate of change occurred lastly in the DTA and MA for all metrics, yet the DTA exhibited the most dramatic changes overall and the MA the least dramatic. That is, of the five segments studied, results tended to be bounded by those in the DTA and MA.

### Intramural stress as a regionally specific initiator of progressive arterial pathology in progeria

Progeria is a mechano-sensitive disease. Values of wall stress tend to increase slightly from P42 to steady state values ~P100 in normal arteries (Murtada et al., 2021a), with the specific magnitude depending on region. Consistent with the aforementioned wall thickening (Figs. 1C, S5F-J), however, there was a consistent marked decrease in mean circumferential stress with age in the ATA, DTA, and CCA in progeria with qualitatively similar but less marked changes in the ICA and MA (Fig. 4C). There was similarly a general reduction in mean axial stress in all aging progeria vessels, but especially in the ATA, DTA, and CCA (Fig. 4D).

Notwithstanding the progressive reduction in wall stress with aging in progeria, values at P42 and P100 were markedly higher in the elastic arteries (ATA, DTA, CCA) than in the muscular arteries (ICA, MA). For example (Tables S1-S5), considering a coordinate-invariant scalar metric of the 2D wall stress (first invariant = *σ_θ_* + *σ_z_*) at P100 and region-specific systolic pressures, values in the DTA, ATA, CCA, MA and ICA were, respectively, 299, 149, 141, 147, and only 42 kPa at P100. Associated values of the circumferential component of stress ordered similarly: 138, 80, 51, 40, and only 21 kPa at P100. With the exception of the MA, these values of wall stress were even higher at P42, noting that the arteriopathy appeared to emerge between P42 and P100. Importantly, this ordering of stress magnitudes reflects well the degree of regional disease severity (Figs. 1A, S9).

### Arterial effects in progeria mirror, but exceed, effects in natural aging

Finally, because progeria is a condition of rapid aging, we compared key functional readouts (biomechanical metrics) in two elastic (ATA and DTA) and one muscular (ICA) artery in progeria at P42 (late maturation) and P168 (near end-of-life) against normal (WT) values in mice naturally aged to P168 (age-matched) or P850 (~2.3 years of age, near end-of-life). Of the eight metrics considered (five shown in Fig. 5), there were greater reductions in elastic energy storage in progeria and much lower biaxial stresses, again most evident in the proximal arteries. Note, too, the general luminal narrowing in progeria in contrast to the slight luminal enlargement in natural aging, a greater thickening of the wall in progeria, and a general lack of circumferential stiffening in progeria. Together, these multiple changes resulted in elevated PWVs that were comparable at end-of-life, at P168 in progeria and P850 in natural aging, in the ATA and ICA, though for different reasons. Importantly, the calculated PWV was yet much greater in progeria in the DTA, which again appeared to exhibit the most severe vascular phenotype (Figs. 1A, S9), particularly near end of life.

**Fig. 5.**
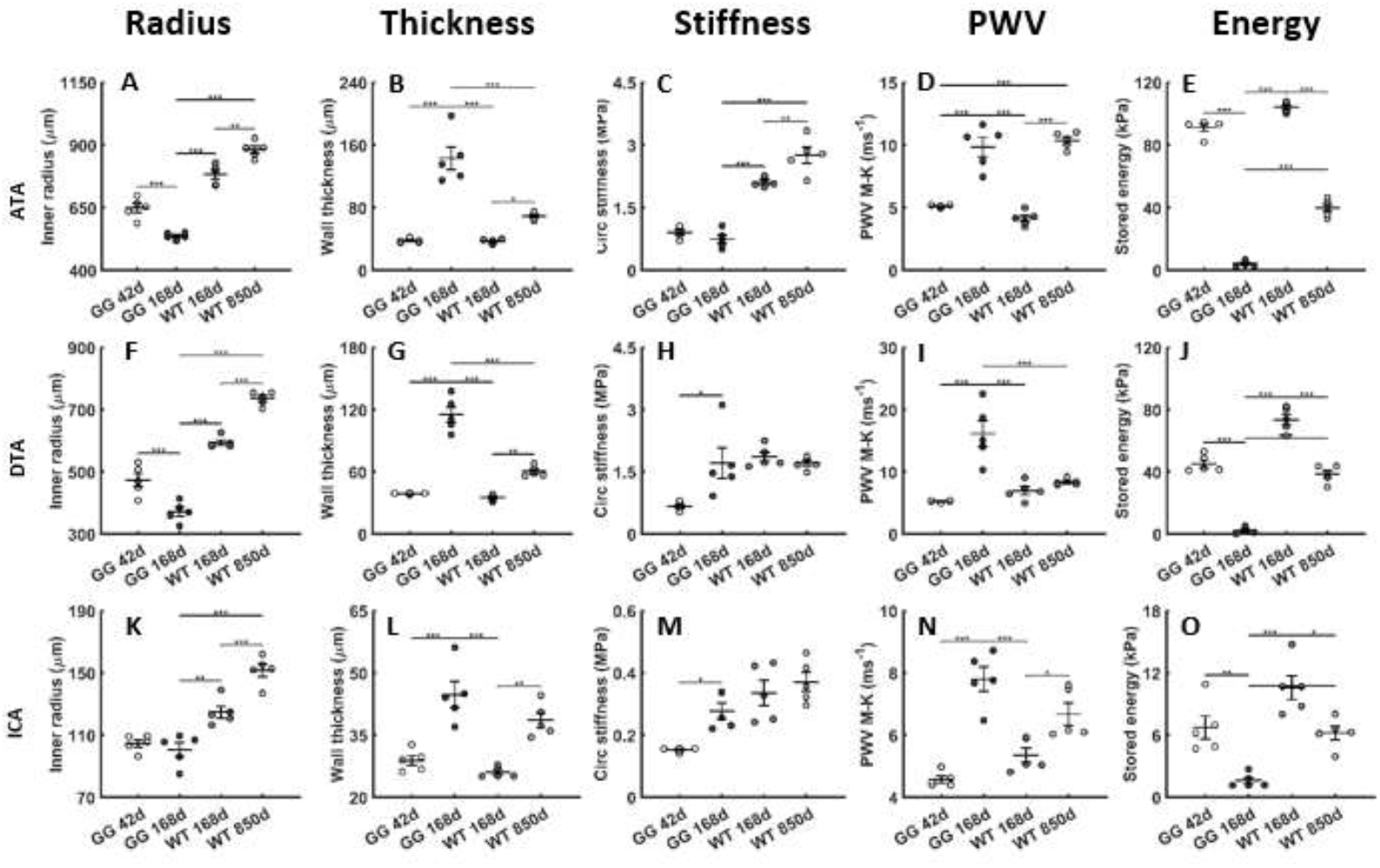
Comparison of key geometric (inner radius, wall thickness) and material/structural (circumferential material stiffness, pulse wave velocity – PWV, and elastically stored energy) metrics for three arteries (ascending thoracic aorta – ATA, descending thoracic aorta – DTA, and internal carotid artery – ICA) at multiple ages: postnatal days P42 and P168 in progeria (GG) and P168 and P850 in wild-type (WT). In general, natural aging results in increased luminal radius, wall thickness, material and especially structural (PWV) stiffness and decreased energy storage. By contrast, progeria associates with luminal narrowing, dramatic wall thickening, marked increases in PWV, and dramatic decreases in energy storage. Note the different scales on the ordinates in some cases, especially for PWV in the DTA, which had the most dramatic phenotype in progeria. *, **, and *** denote statistical differences at *p* < 0.05, 0.01, 0.001.

## DISCUSSION

Considering the thoracic aorta as the archetype of an elastic artery, we and others have reported dramatic evolving changes in geometry, wall composition, material properties, and function during development and maturation in healthy WT mice. The period P7-P14 is one of rapid and dramatic extracellular matrix accumulation (Kelleher et al., 2004), with elastic lamellar structures becoming biomechanically mature by P21 though overall biomechanical properties do not reach steady state mature (homeostatic) values until ~P56 (Le et al., 2015; Murtada et al., 2021a). For example, passive circumferential wall stress at mean arterial pressure is low in the DTA until P10 (≤30 kPa), but then begins to increase rapidly by P21 (~80 kPa) and through P42 (>200 kPa) after which it settles to its normal near steady state level (>250 kPa) by P56. This time-course of increases in stress mirrors that for increases in aortic radius, which are driven largely by the increasing cardiac output that is needed to meet the demand of the increasing body mass of the growing mouse.

We previously showed that the elastic lamellar structures of the medial layer appear normal at P100 in the thoracic and abdominal aorta of *Lmna^G609G/G609G^* progeria mice (Murtada et al., 2020), suggesting that the primary effects of this *Lmna* mutation on aortic extracellular matrix deposition and organization emerge after P21 (when elastic layers normally mature), a period normally characterized by stresses having values increasingly >80 kPa. For this reason, and because these progeria mice begin to show marked losses in body mass ~P42 (Murtada et al., 2020), we selected P42 as a late developmental age of interest, just prior to the aorta reaching biomechanical maturity. The original report of these progeria mice showed 50% survival at P103 despite an absence of cardiac dysfunction (Osorio et al., 2011). By contrast, we showed that placing the chow on the floor of the cage increased their survival to ~P150, with marked left ventricular diastolic dysfunction emerging by ~P140 (Murtada et al., 2020). Since then, we have found that their survival can be extended another ~20%, to ~P180, with 50% survival at P168. This ~63% increase in mean survival (from P103 to P168), achieved by switching from a normal chow to a hydrated soft gel-based chow placed on the floor of the cage, suggests that the mice may have died due to dehydration / malnutrition in the original study (cf. Kreienkamp et al., 2018). Regardless, the present increase in mean survival resulted in the particularly severe aortic phenotype seen at P168 herein (Fig. 1A), including aortic calcification as observed in human progeria patients (Stehbens et al., 1999). We submit that the present late-stage results have greater clinical utility than many prior studies using these mice. In summary, in addition to P42, we chose P100 (prior to diminished cardiac function), P140 (a time of aortic changes consistent with developing diastolic dysfunction), and P168 (the current near end of life period in these mice) as critical ages of cardiovascular interest in *Lmna^G609G/G609G^* mice.

The present histo-mechanical results significantly extend vascular findings reported in prior studies on mouse models of progeria that did not contrast findings in elastic versus muscular arteries or quantify the disease progression (e.g., Varga et al., 2006; Capell et al., 2008; Osorio et al., 2011; Kreienkamp et al., 2018; del Campo et al., 2019; Murtada et al., 2020). Our temporal data reveal complex, differential changes in key geometric and mechanical metrics across five arterial segments. Of note, elastic energy storage appeared similar in progeria to that in WT at P42, except in the DTA (32% lower), but dropped dramatically from P42 to P168 in all five segments and especially in the proximal vessels where elasticity is essential to function. Elastic energy storage in the ATA during isovolumic contraction aids left ventricular diastolic filling by helping to lift the base of the heart during isovolumic relaxation, while elastic energy stored in the ATA and DTA during systole allows the aorta to recoil during diastole and augment antegrade and retrograde flow, including coronary perfusion. Hence, the dramatic progressive loss of energy storage in the proximal aorta is expected to compromise left ventricular diastolic function, consistent with a prior echocardiographic observation at P140 in these mice (Murtada et al., 2020).

Progeria is a highly mechano-sensitive condition (Boers et al., 2004; Verstraeten et al., 2008), with high extracellular stresses causing nuclear damage and cell death in *in vitro* studies (Kim et al., 2021). For this reason (cf. Chambliss et al., 2013), disruption of the LINC complex can lessen, though not eliminate, the disease phenotype in progeria (Kim et al., 2018), apparently by reducing the mechanical stress transmitted to the nucleus. Nevertheless, it has been difficult to estimate *in vivo* ranges of extracellular stresses that prevent or promote this damage. Given the effective organization of elastic lamellar structures in the progeria aorta (which occur by P21 when circumferential stress is ≤80 kPa) as well as the differential stresses and arterial phenotypes observed across all segments, with the most severe phenotype in the aorta where stress is highest (normally >250 kPa in maturity) and the least severe phenotype in the least stressed internal carotid and mesenteric arteries (normally <70 kPa in maturity), it appears that circumferential stresses <80 kPa may be tolerated by mural vascular cells in progeria. By contrast, mean intramural stresses were ~150 kPa in the DTA at both P42 and P100, well in excess of values at P21 or in the muscular arteries, apparently contributing to the dramatic deterioration in its biomechanical phenotype.

Because the DTA exhibited the greatest decline in this mouse model of progeria (Fig. 1A), and because the media is the most highly stressed layer in the normal aortic wall (Latorre et al., 2021), we focused primarily on the aorta and in particular the SMCs to identify underlying mechanistic time-courses. Together bulk RNAseq (at P120) and scRNA-seq (at P100, P140, and P168) revealed marked deviations in aortic cell phenotype in progeria, suggesting switches from the normal contractile-synthetic-degradative SMC phenotype that exists in WT in maturity (Estrada et al., 2021) to a more synthetic phenotype (P42-P100), then pro-apopotic phenotype (by P100-P120), then chondrogenic phenotype (by P120-P140), and finally osteogenic (by P140-P168) phenotypes in progeria. The mid-stage disease progression in the aorta is consistent with a mechanical stress-mediated damage to DNA (Liu et al., 2013) and associated diffuse cell death (previously shown by P140, but not at P100 when contractility was lessened; Murtada et al., 2020) that is followed by an excessive accumulation of proteoglycans (previously seen by P140, but not at P100, to be primarily aggrecan; Murtada et al., 2020) and late-stage calcification (seen histologically at P168, not P140, herein; Figs. 1, S4). Moreover, such a progression is consistent with prior non-longitudinal reports near end-of-life in humans and mice that reported SMC loss and accumulations of proteoglycans and calcification in progeria (Stehbens et al., 2001; Varga et al., 2006; Olive et al., 2010). Importantly, vascular SMCs can exhibit chondrogenic (indicated, in part, by Sox9, Acan, Col2a1) and osteogenic (indicated, in part, by Runx2, Spp1) phenotypes in natural aging, also often associated with SMC apoptosis (Johnson et al., 2006; Durham et al., 2018). The present data show a peak in expression of the pro-apoptotic transcript Bax at P100 (suggesting a delay in actual cell death), which decreased thereafter though yielding a dramatic loss of SMCs seen at P168 on multiphoton imaging (Fig. 2) that was confirmed by TUNEL staining (Fig. 1). By contrast, the chondrogenic marker Sox 9 peaked at P140 and remained elevated while the osteogenic marker Runx2 (aka Cbfa1) peaked at P140 and dropped slightly thereafter (again suggesting a delay in accumulation, seen mainly at P168).

There are many possible reasons for an association of apoptosis with an osteochondrogenic phenotype in neighboring remnant cells, including loss of vital cell-cell and cell-matrix connections, a reduction in anti-calcific signals from the loss of healthy cells (e.g., reduced matrix γ-carboxyglutamic acid-rich protein), and even release of calcium stores by dead cells into the extracellular matrix, which can help nucleate calcium-rich deposits (Clarke et al., 2008; Sage et al., 2010; Briot et al., 2014; Ngai et al., 2018). Overproduction of glycosaminoglycans within the aorta can also contribute to increased calcification (Purnomo et al., 2013). Regardless, consistent with the observed increase in the aggrecan transcript (Acan) herein, the accumulated proteoglycans (on Movat staining) were often excessive, even relative to that seen in the murine aorta in natural aging up to two years (Ferruzzi et al., 2018). Such accumulations can adversely affect wall function in many ways, including compromising the cellular mechano-sensing that is needed to mechano-regulate the extracellular matrix (Roccabianca et al., 2014; Humphrey et al., 2014), though herein the accumulated proteoglycans also contributed at the tissue-level to the measured decreases in tensile material stiffness and increases in wall thickness, the latter which increased PWV, which contributes to diastolic dysfunction (Desai et al., 2009; Townsend et al., 2015), the most common diagnosis in children in a clinical study of progeria (Prakash et al., 2018). Similarly, the late calcification, seen only in the larger arteries, increased wall stiffness further, and thus PWV.

Our biomechanical findings in combination with histological and transcriptomic data thus suggest a new hypothesis for progressive preferential damage in segments of the systemic vasculature: arterial mechanical stresses greater than ~80 kPa drive nuclear damage and thus SMC death in progeria, leaving a remnant cohort of SMCs to develop an osteochondrogenic phenotype that drives the accumulation of mural proteoglycans that contributes to wall thickening and increases PWV, with late calcification exacerbating this situation in the most affected regions (Fig. S10). This excessive accumulation of proteoglycans also changes the extracellular milieu in which the remnant cells reside – biomechanically, the marked decreases in tensile stress (e.g., from >250 kPa to ~40 kPa at P168 in the DTA) can exacerbate apoptosis (Bayer et al., 1999) despite increases in hydrostatic stress due to Gibbs-Donnan swelling (Roccabianca et al., 2014) and biochemically, the significant changes in ligand presentation to the cells can affect cell phenotype and even cell survival (Michel, 2003; Sazonova et al., 2015). Toward this end, it is not clear if the late reduction in intramural stress (40-60 kPa in the progeriod DTA at P140 and P168) was protective against the nuclear vulnerability that arises in progeria due to loss of functional lamin-A and enabled the remnant osteochondrogenic cells to survive, or if it drove reductions in mechano-regulated homeostatic mechanisms, further phenotypic modulation, and even increased apoptosis. There is need for additional studies to focus on these possibilities. Although we suggest that the very different values of wall stress along the arterial tree can either hasten (in the aorta) or delay (in muscular arteries) stress-mediated nuclear damage in progeria-vulnerable cells, differential remodeling between the DTA and MA has similarly been observed in angiotensin-II induced hypertension in WT mice (Murtada et al., 2021b). There is also a need for additional studies to investigate whether the present differential findings depend in part on intrinsic differences in the remodeling capability of elastic and muscular arteries or if the initially lower intramural stresses in the muscular arteries simply resulted in less damage to the vulnerable cells, which appears likely in progeria. Another possibility is that SMCs tend to have a mechano-sensitive maintenance phenotype (contractile-synthetic-degradative) in elastic arteries but a more vasoactive regulatory (contractile) phenotype in muscular arteries, and progeria is a mechano-sensitive condition. We did not focus on other cell types, which yet deserve attention. The proteoglycans and calcification also presented in the adventitial layer but we did not examine fibroblasts, which at least in skin drive aggrecan production in progeria (Lemire et al., 2006). We observed that increases in Runx2 and Spp1 transcripts emerged not only in SMC clusters, but also in inflammatory cells in progeria. Macrophages are well known to be involved in vascular calcification (Tintut et al., 2002; Li et al., 2020), and deserve more attention in progeria (Benedicto et al., 2021). Finally, progerin expression in endothelial cells contributes to the cardiovascular phenotype (Osmanagic-Myers et al., 2019), but these cells were not studied herein beyond noting a heterogeneous reduction in endothelial cell density at P168 in the DTA via multiphoton microscopy. Much remains to be studied.

Although the reduced luminal caliber seen in progeria may have resulted from inward remodeling, it may have been an allometrically appropriate response to reduced cardiac output given the decreased body mass after P42 (Murtada et al., 2020). Regardless, this reduction combined with changes in cell-mediated wall stiffness and especially wall thickness to increase the calculated PWV in the central vessels, especially the DTA. Integrated measurements of local PWV are thus recommended as an important functional readout, particularly given reports of increases in carotid-femoral and brachial-ankle PWV in progeria patients (Gerhard-Herman et al., 2012). Given that intramural stress is generally low during normal elastic lamellar development, which is complete in mice near P21, it appears that investigative pharmacological interventions could be started at or after P21 in mice (the time of weaning), which is important since some proposed drugs (e.g., mTOR inhibitors; Dubose et al., 2018) can adversely affect somatic growth. Finally, given that the DTA exhibited the most severe compositional and functional phenotype, assessments should focus on or at least include this critical segment of the central vasculature to evaluate interventional efficacy.

In summary, time-course data herein suggest that mechanical stress-induced SMC death drives an osteochondrogenic phenotypic modulation of remnant neighboring SMCs and initiates differential disease in systemic arteries in progeria, with highly stressed elastic arteries affected the earliest and most severely. The early loss of elastic energy storage in the proximal aorta coupled with the progressive increase in PWV likely combine to adversely affect left ventricular diastolic function, the most commonly diagnosed condition in progeria patients (Prakash et al., 2018). Because lamin-A appears essential for stress shielding the nucleus from high mechanical stresses transmitted from the extracellular matrix, thus ensuring appropriate nuclear mechanotransduction, there appears to be a need both to preserve functional lamin-A and to clear its aberrant form, that is, progerin (Gonzalez-Gomez et al., 2022). Mitigating smooth muscle cell loss and severe phenotypic modulation promise to have important functional implications in progeria patients.

## EXPERIMENTAL METHODS

### Animals

All live animal procedures were approved by the Yale IACUC. *Lmna^G609G/G609G^* (progeria) mice were generated by breeding *Lmna^G609G/+^* mice, which yielded wild-type *Lmna^+/+^* (WT) mice as littermate controls, noting that these mice are on a C57BL/6 background. Male and female mice were euthanized with an intraperitoneal injection of Beuthanasia-D at the appropriate postnatal age (P42, P100, P140, or P168), and five arterial segments (ATA, DTA, CCA, ICA, and MA) were gently excised and prepared for biomechanical phenotyping, with a sixth segment (abdominal aorta) included for RNA sequencing, noting that changes in the thoracic and abdominal aorta are comparable in progeria and independent of sex (Murtada et al., 2020). We thus report mixed-sex findings herein. In all cases, death was confirmed by loss of cardiovascular function following exsanguination. Finally, ATA, DTA, and ICA segments were similarly excised from separate C57BL/6J WT mice at P168 and P850 (~2.3 years of natural aging) for comparison.

### Mechanical Testing

Using a custom computer-controlled experimental system and previously established protocols (Murtada et al., 2020), vessels were placed within a standard Hank’s buffered physiologic solution at room temperature to minimize smooth muscle contractility and thereby to focus on passive biomechanical properties. Vessels were then preconditioned via four cycles of pressurization while length was held fixed at specimen-specific *in vivo* values. Next the vessels were subjected to three pressure-diameter (*P* − *d*) protocols, with luminal pressure cycled from 10 to a region-specific maximum value (140 mmHg for ATA, DTA, and CCA, and 90 mmHg for ICA and MA) while axial stretch was maintained fixed at either the specimen-specific *in vivo* value or ±5% of this value. Finally, four axial force-length (*f* − *l*) protocols, with force cycled between 0 and a value equal to its maximum measured during the pressurization test at 5% above the *in vivo* axial stretch, with luminal pressure maintained fixed at {10, 60, 100, or 140 mmHg} for the ATA, DTA, and CCA, and {10, 30, 60, 90 mmHg} for the ICA and MA (Murtada et al., 2021b). Distending pressure, applied axial force, outer diameter, and axial length were recorded on-line for all seven cyclic protocols. Wall thickness was measured in the unloaded configuration using a dissecting microscope, which under the assumption of incompressibility allowed wall thickness to be calculated at each pressure-force state.

### Constitutive Modeling

Multiple consistently calculated metrics collectively enable detailed biomechanical phenotyping of arteries across vessel types, sex, age, and genotype. These metrics include intramural biaxial stress and stretch, material stiffness, and elastic energy storage as well calculated local pulse wave velocity (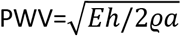 where *E* is a linearized material stiffness, *h* wall thickness, *a* inner radius, and *ϱ* the mass density of the fluid). We used an independently validated (Schroder et al., 2018) four-fiber family constitutive relation to quantify the passive behavior (Ferruzzi et al., 2018; Murtada et al., 2020), which can be written in terms of the stored energy *W* as,

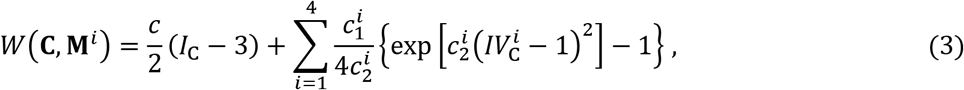

where *c* (kPa), 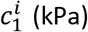, and 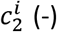 are material parameters (*i* = 1,2,3,4 denote four predominant fiber family directions), which are determined via nonlinear regression of *P* − *d* and *f* − *l* data from the last cycle of unloading in each of the seven protocols. Unloading data enable calculation of the non-dissipated (elastic) energy available to work on the luminal fluid. Finally, *I*_C_ = *tr***C** and 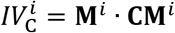 are coordinate invariant measures of the finite deformation, with the right Cauchy-Green tensor **C** = **F**^T^**F** computed from the deformation gradient tensor **F** = diag[*λ*_*r*_, *λ*_*θ*_, *λ*_*z*_], with *det***F** = 1 because of assumed incompressibility. The direction of the *i*^*th*^ family of fibers is defined by 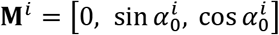, with 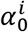 denoting a fiber angle relative to the axial direction in the traction-free reference configuration. Based on prior microstructural observations from multiphoton microscopy, and the yet unquantified effects of cross-links amongst the multiple families of fibers, the four predominant families were: axial 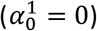, circumferential 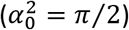, and symmetric diagonal 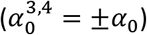. The value of *α*_0_ was included among the eight model parameters determined via the nonlinear regression. Values of mean biaxial stress and stiffness were computed from the stored energy function and calculated at individually measured values of pressure and at a common pressure.

### Time courses

Motivated by prior findings in aortic development (Murtada et al., 2021a), we used a simple relation to describe time-dependent changes in the various metrics, denoted generically as *ξ*. That is, we computed best-fit values of four model parameters (*A*, *B*, *C*, *D*) in the relation

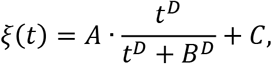

where *t* is time, which was found to fit each time-course data well. The maximum rate of change, namely *dξ*/*dt*, was determined easily, and so too the time at which this value occurred. Finally, we also used this relation to find the time at which the value of *ξ* reached 90% of its near end of life value, *ξ*(*P*168).

### Multiphoton Microscopy

Following mechanical testing, DTAs from P168 *Lmna^G609G/G609G^* and wild-type *Lmna^+/+^* mice were subjected to *ex vivo* biaxial loading conditions of 100 mmHg luminal pressure and sample-specific *in vivo* axial stretch. Two-photon microscopy yielded 3-D images of samples under physiological loading throughout the two primary (medial and adventitial) layers of the wall. The LaVision BioTec TriMScope had a Titanium-Sapphire laser tuned at 820 nm and a 20x objective lens for water immersion (N.A. 0.95). Separated signals of second harmonic generation from fibrillar collagen (wavelength: 390-425 nm), two-photon auto-fluorescence from elastin (500-550 nm), and fluorescence from cell nuclei (stained with cell-permeant nucleic acid SYTO red fluorescent stain for 2 hours, above 550 nm) were obtained simultaneously. Regions of interest of 500μm x 500 μm x 100 μm (in axial, circumferential and radial axes, respectively) were imaged twice (dorsal and ventral sides) within the mid-length of the aorta while maintaining the axial stretch and 100 mmHg pressure. Multiple microstructural parameters were obtained from the 3-D images: wall thickness and relative adventitia:media ratio; volume fractions of wall components (i.e., collagen, elastin and cell nuclei) in the adventitial and medial layers; in-plane (i.e., circumferential-axial) collagen architecture quantified as fiber straightness, bundle width, and orientation distribution (comprising preferred absolute fiber orientations and concentrations that quantify alignment along the preferred orientation); density of cell nuclei per volume of adventitia and media, and density of endothelial cell nuclei per area of luminal surface. Further details are found elsewhere (Cavinato et al. 2021).

### Histology

Following mechanical testing, vessels were fixed in their unloaded state in a 10% formalin solution for 24 h, then placed in a 70% ethanol solution, embedded in paraffin, sectioned (5 μm thickness), mounted, and stained using one of three standard stains: Movat’s pentachrome (Mov), which stains elastin and nuclei black, collagen fibers yellowish, glycosaminoglycans blue, and cell cytoplasm red; picrosirius red (PSR), which reveals fibrillar collagens colorimetrically (red-orange, yellow-green) under polarized light; Alizarin Red, which stains calcium red. Finally, an immuno-histochemical TUNEL assay identified apoptotic cells (brown for cell nuclei having DNA fragmentation). Images were acquired using an Olympus BX/51 microscope equipped with a DP70 digital camera (effective sensor resolution of 4080 x 3072 pixels, corresponding to a pixel size of 2.1 μm at a 2/3” sensor size) and using a 20x objective (UPlanFl 20x, NA 0.50, optical resolution at λ = 400 nm at ~0.49 μm) and 0.5x tube lens, resulting in 10x magnification and hence an image resolution of 0.21 μm (fulfilling the Nyquist criterion). Images were recorded using Olympus CellSens Dimension software. When arterial cross-sections exceeded the field-of-view, multiple images were acquired and stitched using Image Composite Editor software (Microsoft Research).

### Bulk RNA Sequencing and Analysis

Crushed whole abdominal aortic segments were immersed in RLT lysis buffer (Qiagen N.V., Venlo, Netherlands) and vortexed. Total RNA was isolated using an RNeasy Mini Kit and DNase Digestion Set (Qiagen) according to the manufacturer’s protocol. Next-generation, whole-transcriptome sequencing was performed using a NovaSeq 6000 System (Illumina, Inc., San Diego, CA) at the Yale Center for Genome Analysis. RNA-Seq reads were aligned to a reference genome (GRCm38/mm9) with Gencode annotation using HISAT2 for alignment, and StringTie for transcript abundance estimation. Differentially expressed genes (DEGs) were obtained from raw gene counts using DESeq2 v1.26.0 in R v3.6.2. Benjamini-Hochberg correction as implemented in DESeq2 was used to obtain *p*-values adjusted for multiple comparisons (*p*_adj_); genes with *p*_adj_ ≤ 0.05 were assumed to be DEGs. To reduce spurious large fold changes for genes with low read counts or high coefficients of variation, log_2_ fold change was performed using the apeglm method. DEGs were used for gene ontology (GO) analysis for biological processes using goseq v1.38.0, run separately for up- and down-regulated genes. Benjamini-Hochberg correction again adjusted for multiple comparisons; categories with *p*_adj_ ≤ 0.05 were considered significant. Additional pathway and ontology analyses were performed using enrichR 2.1 and the Molecular Signatures Database (MSigDB) hallmark gene set collection. Further details can be found elsewhere (Li et al., 2020).

### Single-cell RNA sequencing

Whole aortas (thoracic + abdominal) were excised, rinsed in cold PBS, opened, and sliced into small fragments. The minced tissue was incubated in 0.5 ml DMEM with 10% FBS, 1.5 mg/ml collagenase A, and 0.5 mg/ml elastase for 3 hours at 4°C while the tissue fragments were gently titrated using a pipette every 15 min. Cells were isolated using a 40 μm filter and incubated with cell-impermeant viability dye (Thermo Fisher) for 20 min at 4°C. Viable cells were processed for scRNA-seq library preparation using the Chromium™ Single Cell Platform (10x Genomics) per the manufacturer’s protocol. Briefly, single cells were partitioned into Gel Beads in Emulsion using the 10x Genomics system, followed by cell lysis and barcoded reverse transcription of RNA, cDNA amplification, and shearing, with 5’ adaptor and sample index attachment. The scRNA-seq libraries were sequenced on a HiSeq 4000 System (Illumina) at the Yale Center for Genome Analysis.

Data from the barcoded library sequence were processed through Cell Ranger (10x Genomics) and this output was further processed in R using Seurat. The reference mouse genome was customized by the addition of a reference genome sequence. The data were filtered; cells with fewer than 200 genes or 250 RNA features or more than 10% expression of mitochondrial genes were eliminated. The read counts were normalized in Seurat using SCTransform. UMAP projections were used for visualization. Further details can be found elsewhere (Li et al., 2020).

### Statistics

Most data are presented as means±standard errors and analyzed using one- or two-way analysis of variance (ANOVA) as appropriate, with post-hoc Bonferroni tests; *p* <0.05 was considered significant. DEG was determined by bulk RNA-sequencing and corrected for multiple comparisons using the Benjamini-Hochberg method. All mechanics and histology data processing and statistics was performed in MATLAB R2019a (Mathworks, Natick, MA).

## ACKNOWLEDGMENTS

This work was supported by a grant from the US NIH (R01 HL105297, JDH) and the Masason Foundation (YK).

## CONFLICTS OF INTEREST

The authors declare that they have no conflicts of interest.

## AUTHOR CONTRIBUTIONS

Design of the Project (SIM, YK, GT, JDH), Data Collection (SIM, YK, CC, MW, ABR, BS), Data Analysis and Interpretation (SIM, YK, CC, GT, JDH). Writing and Editing (SIM, YK, CC, GT, JDH).

## DATA AVAILABILITY

All data needed to evaluate the conclusions of the paper are in the paper and the Supplementary Materials. Additional information is available from the corresponding author upon reasonable request.

## ETHICS STATEMENT

All live animal procedures were conducted in accordance with Federal Regulations and were approved by the Yale Institutional Animal Care and Use Committee prior to beginning the work.

## Supplemental Materials

Figures A1-S9, Tables S1-S6

## Notes

### Competing Interest Statement

The authors have declared no competing interest.

