## Supplemental Materials for "Smooth Muscle Cell Death Drives an Osteochondrogenic Phenotype and Severe Proximal Vascular Disease in Progeria"

**Supplemental Figures**

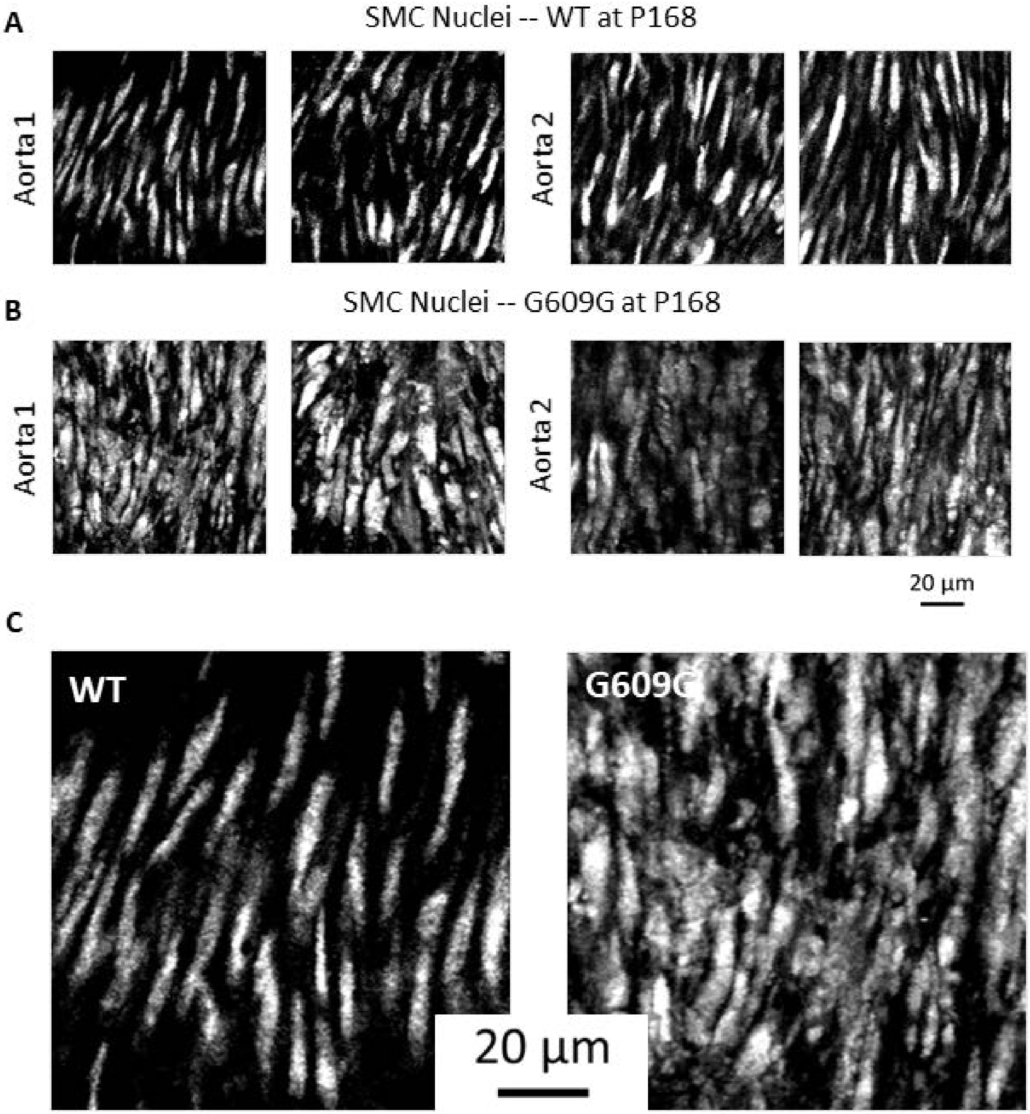


Figure S1. Focus on smooth muscle cell nuclei at postnatal day P168 within the media in two representative (A) wild-type (WT) and (B) progeria (G609G) aortas, with (C) a single representative image for each at a higher magnification. Note the diffuse, distorted shapes in progeria relative to the more uniform spindle shapes in wild-type.

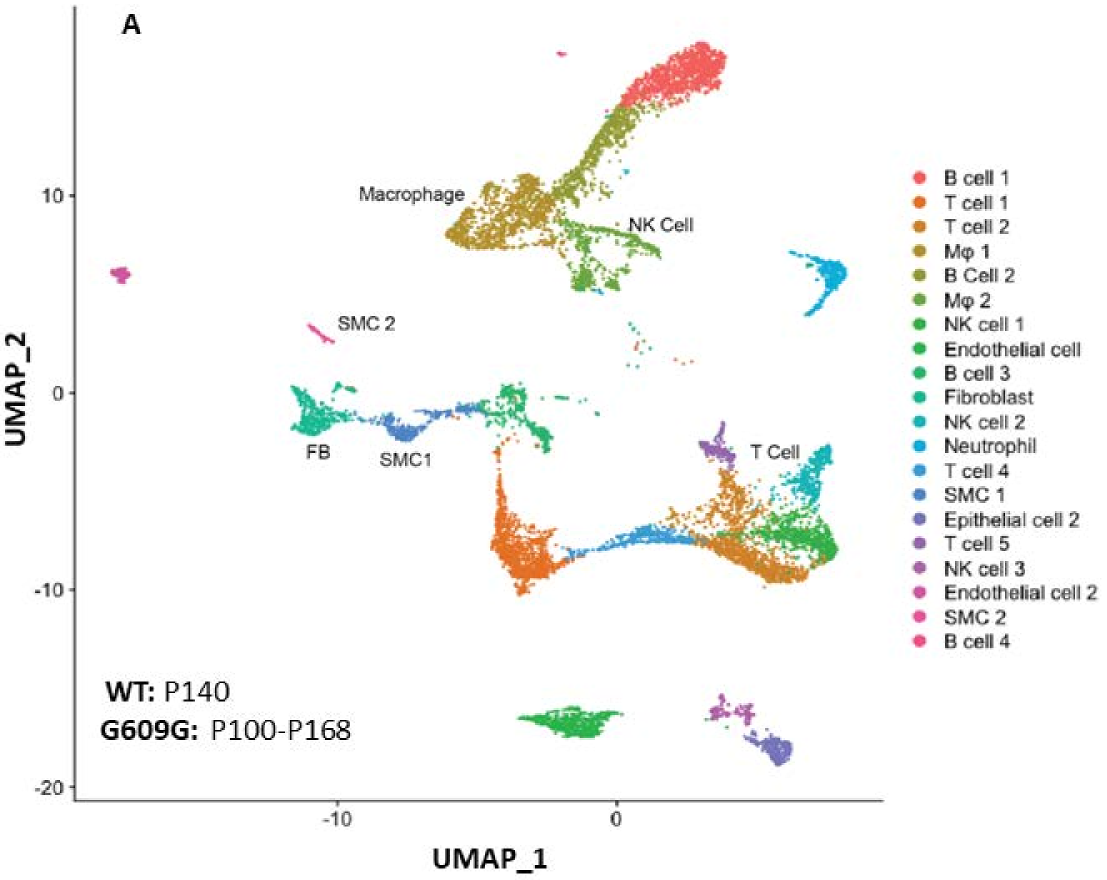


Figure S2. Single-cell RNA sequencing data (scRNA-seq) for the whole aorta (thoracic + abdominal) excised from wild-type (WT) and progeria (G609G) mice at multiple postnatal days from P100-P168. Note the 20 different clusters, including two smooth muscle cell (SMC) clusters, one to be distinguished from a neighboring fibroblast (FB) cluster.

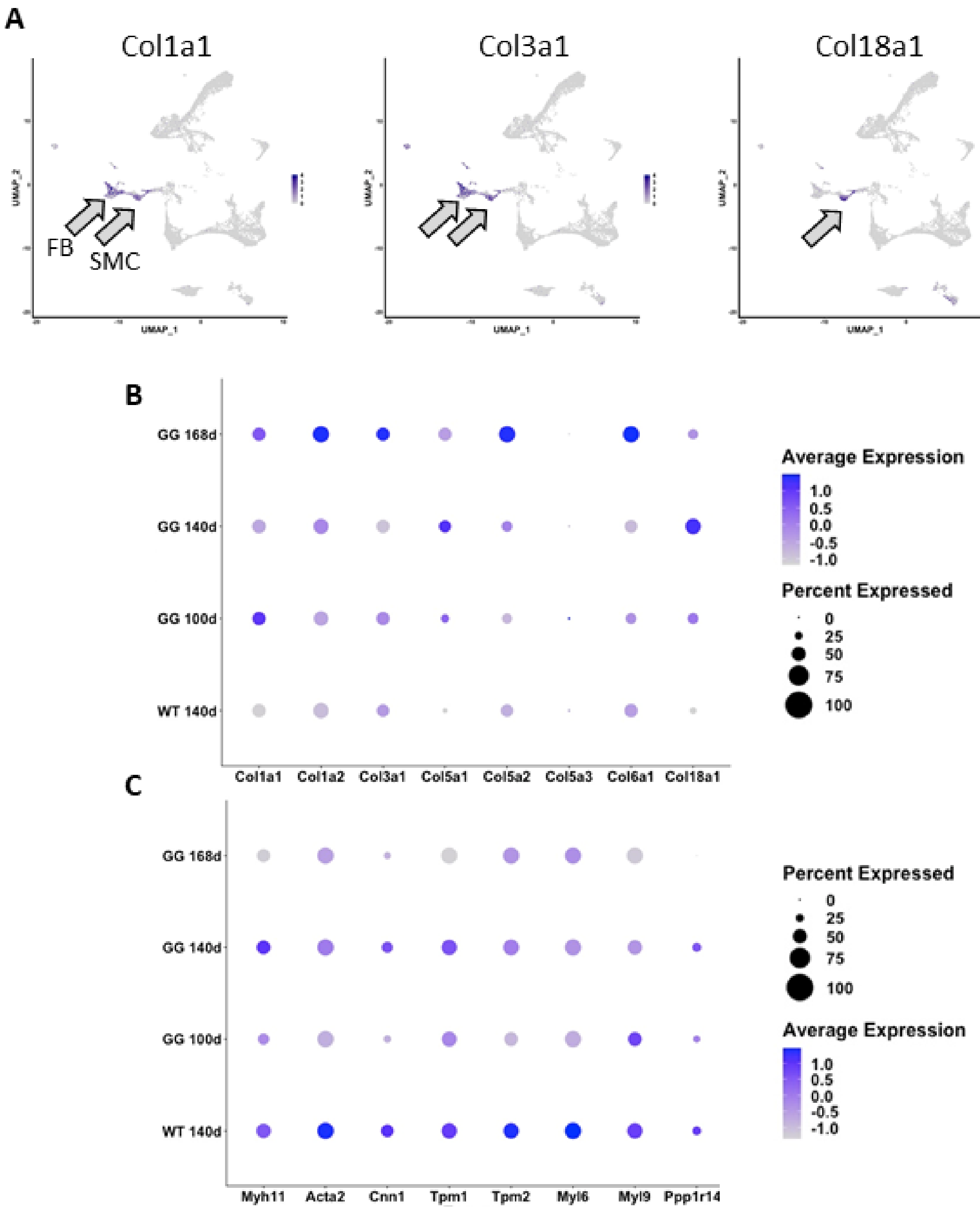


Figure S3. Similar to Figure 3 in the main text, except for scRNA-seq results focusing on (A,B) collagens, primarily for three genes (collagens I, III, XVIII), expressed predominantly within the clusters for fibroblasts and smooth muscle cells, noting that Col18a1 is seen exclusively in a smooth muscle cell cluster. (B,C) Shown, too, are longitudingal data for synthetic and contractile markers in the smooth muscle cells for progeria (GG at 100, 140, and 168 days of age) relative to wild-type (WT) at 140 days of age. The smooth muscle cells in the progeria aorta exhibited a general decrease in contractile markers and increase in synthetic markers with aging.

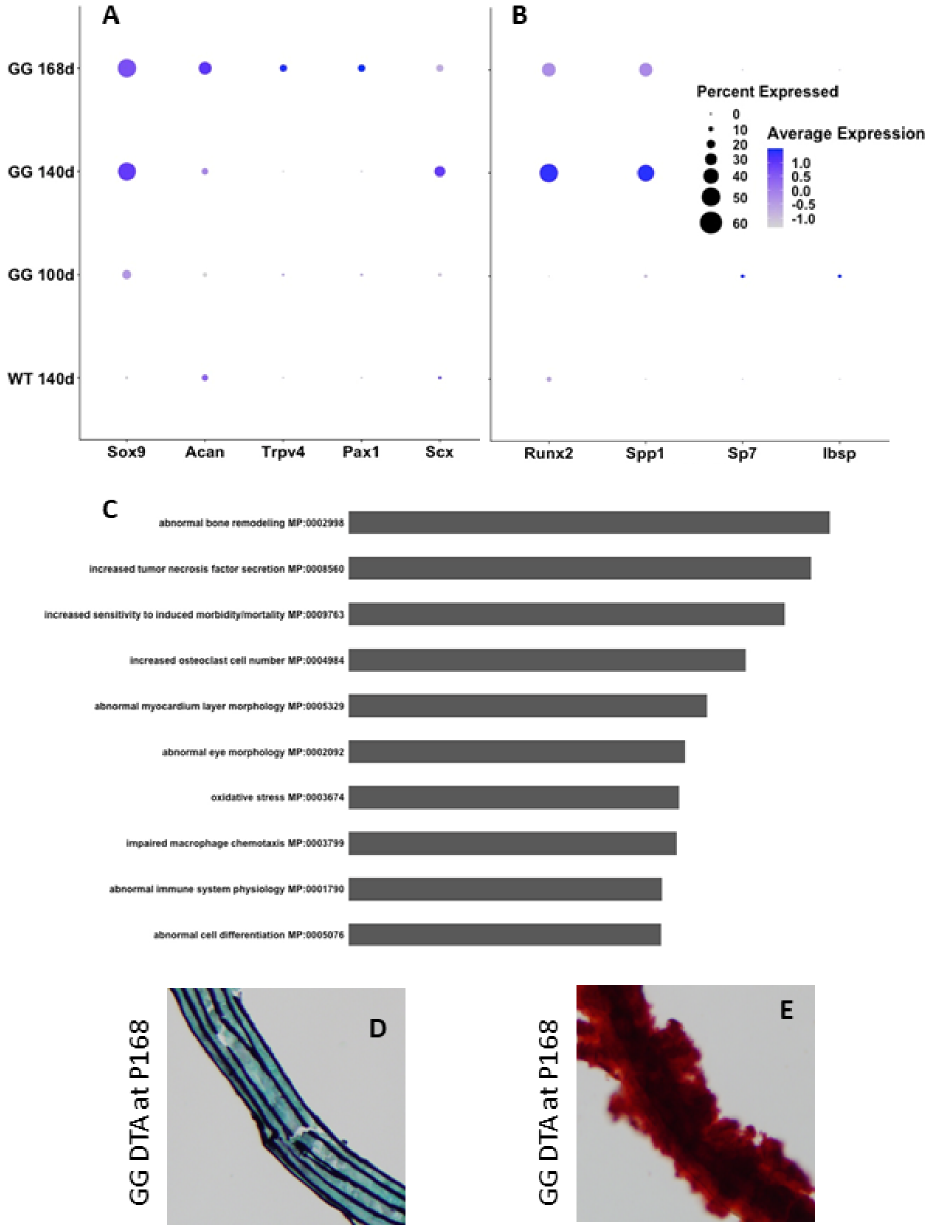


Figure S4. Additional results from scRNA-seq of smooth muscle cells from the aorta, with an associated gene ontology study and confirmatory histology. (A,B) Time-course of results for osteochondrogenic transcripts of interest that emerged in the bulk RNA sequencing (cf. Figure 2A, main text) but shown here for smooth muscle cells: comparison of wild-type (WT) at postnatal day 140d versus progeria (GG) at days 100d, 140d, and 168d. Selected results from a gene ontology analysis that revealed a “bone formation” phenotype in progeria smooth muscle cells. (D,E) Illustrative images from histological sections showing dramatic accumulations of proteoglycans (blue – Movat stain, left) and calcium (red – Alizarin red stain, right) at day 168 in the progeria aorta, consistent with transcriptional modulation. Note that the histological sections of the aorta at P168 were often brittle, easily fragmented. Comparing panel D here with that in Figure 1B (MOVAT P168) shows considerable variation from specimen-to-specimen, but the general accumulation of proteoglycans and calcium in the near end of life thoracic aorta.

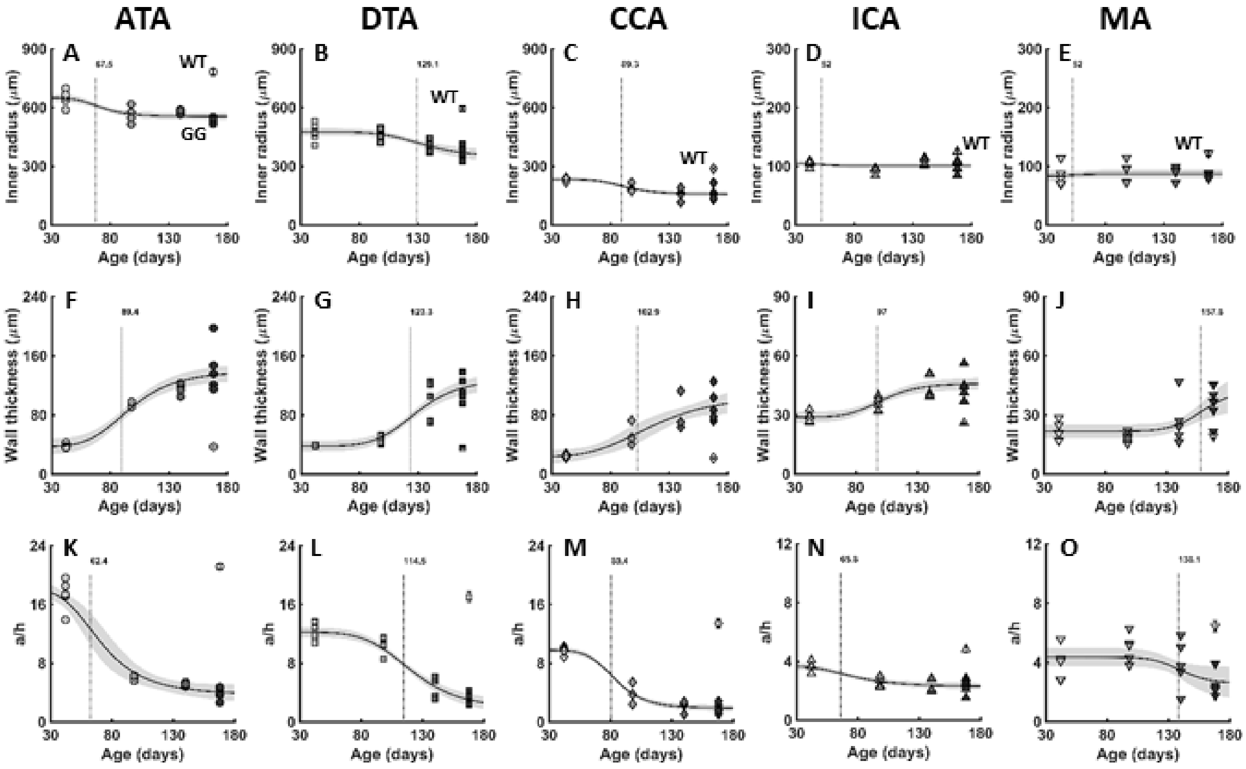


Figure S5. Time-course of changes (postnatal days 42, 100, 140, 168) in three key geometric metrics in progeria for all five systemic vessels of interest: the ascending (ATA) and descending (DTA) thoracic aorta, the common (CCA) and internal (ICA) carotid artery, and the $2^{nd}$ branch mesenteric artery (MA). (A-E) pressurized inner radius $a$, (F-J) pressurized wall thickness $h$, and (K-O) their ratio $a/h$, which is particuarly important in the calculation of both mean circumferential intramural stress (via the equation of Laplace) and pulse wave velocity (via Moens-Korteweg), both of which are shown in Figure 4 in the main text. Note differences in the scale for the quantity of interest (ordinate) across regions. Note, too, that open symbols show, for reference, the wild-type (WT) values at P168.

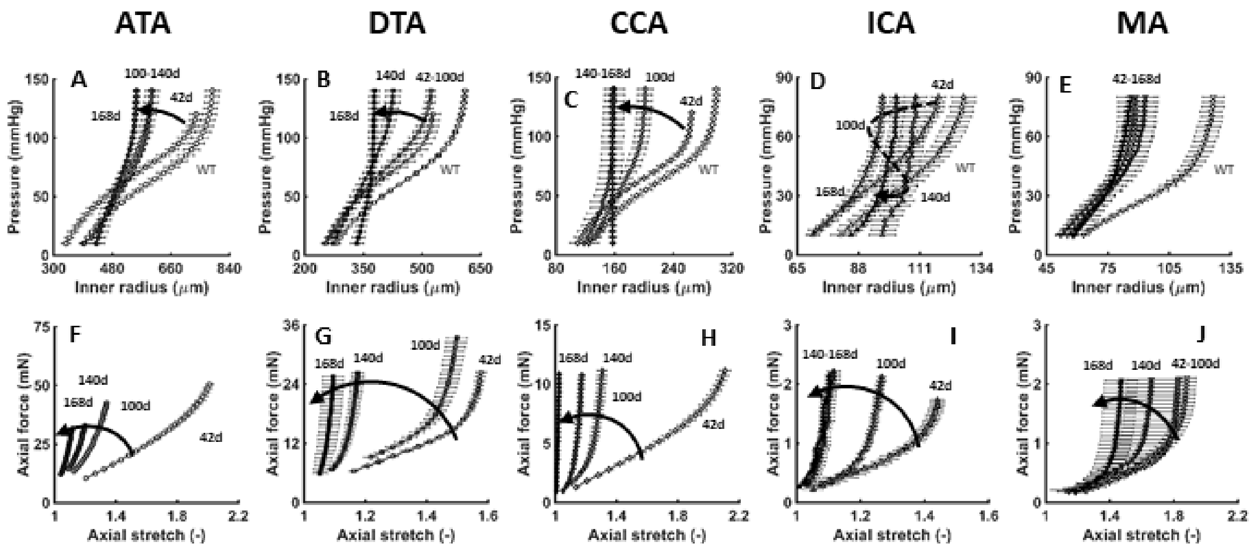


Figure S6. Time-course of changes (postnatal days 42, 100, 140, 168) in circumferential and axial *structural* properties in progeria for all five systemic vessels of interest, the former represented by (A-E) pressure-diameter responses at specimen-specific fixed values of the *in vivo* value of axial stretch and the latter by (F-J) axial force-extension responses at group-specific fixed values of distending pressure. Note differences in the scale for the quantity of interest (ordinate) across regions. Note, too, that open symbols show, for reference, the wild-type (WT) values at P168.

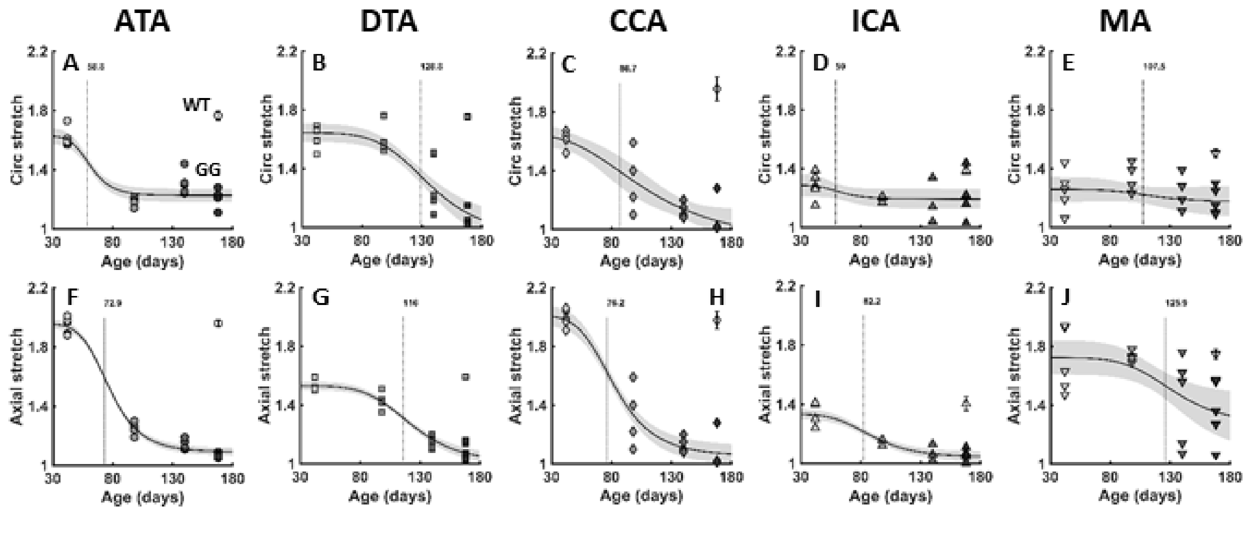


Figure S7. Similar to Figure S5, except for *in vivo* relevant values of (A-E) circumferential and (F-J) axial stretch as a function of postnal day (42, 100, 140, 168) and region (ATA, DTA, CCA, ICA, MA) for progeria mice. Note that open symbols show, for reference, the wild-type (WT) values at P168. See Figure 4 in the main text for associated biaxial intramural stresses.

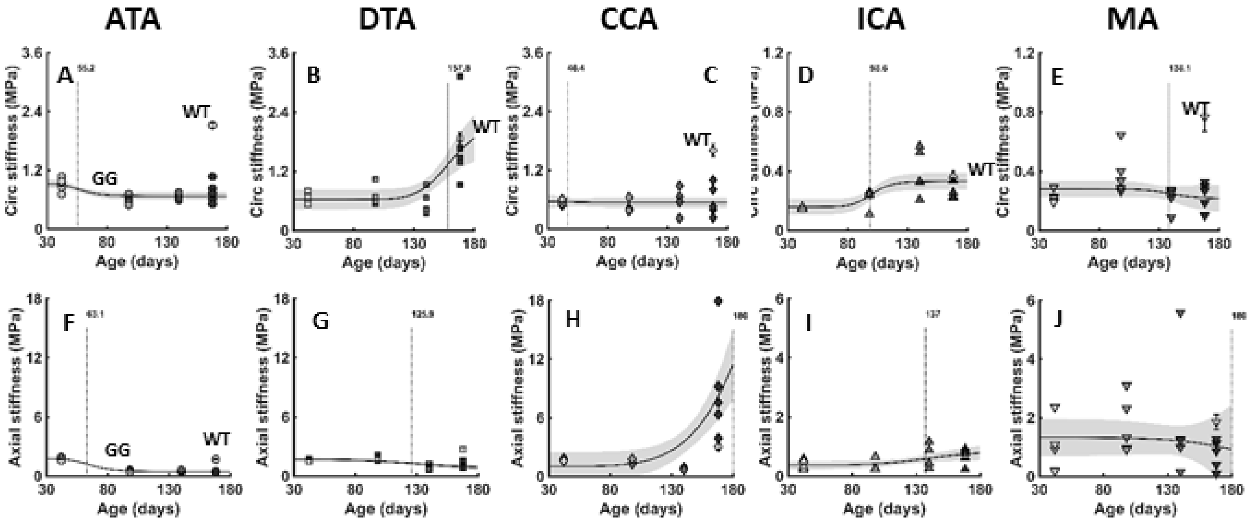


Figure S8. Similar to Figure S7, except for the time-course of changes (postnatal days 42, 100, 140, 168) in circumferential and axial *material* properties in progeria for all five systemic vessels of interest, the former represented by (A-E) circumferential material stiffness and the latter by (F-J) axial material stiffness, both of which are computed from the experimentally determined stored energy function using the theory of small deformations superimposed on large deformations. Note differences in the scale for the quantity of interest (ordinate) across regions. Note, too, that open symbols show, for reference, the wild-type (WT) values at P168.

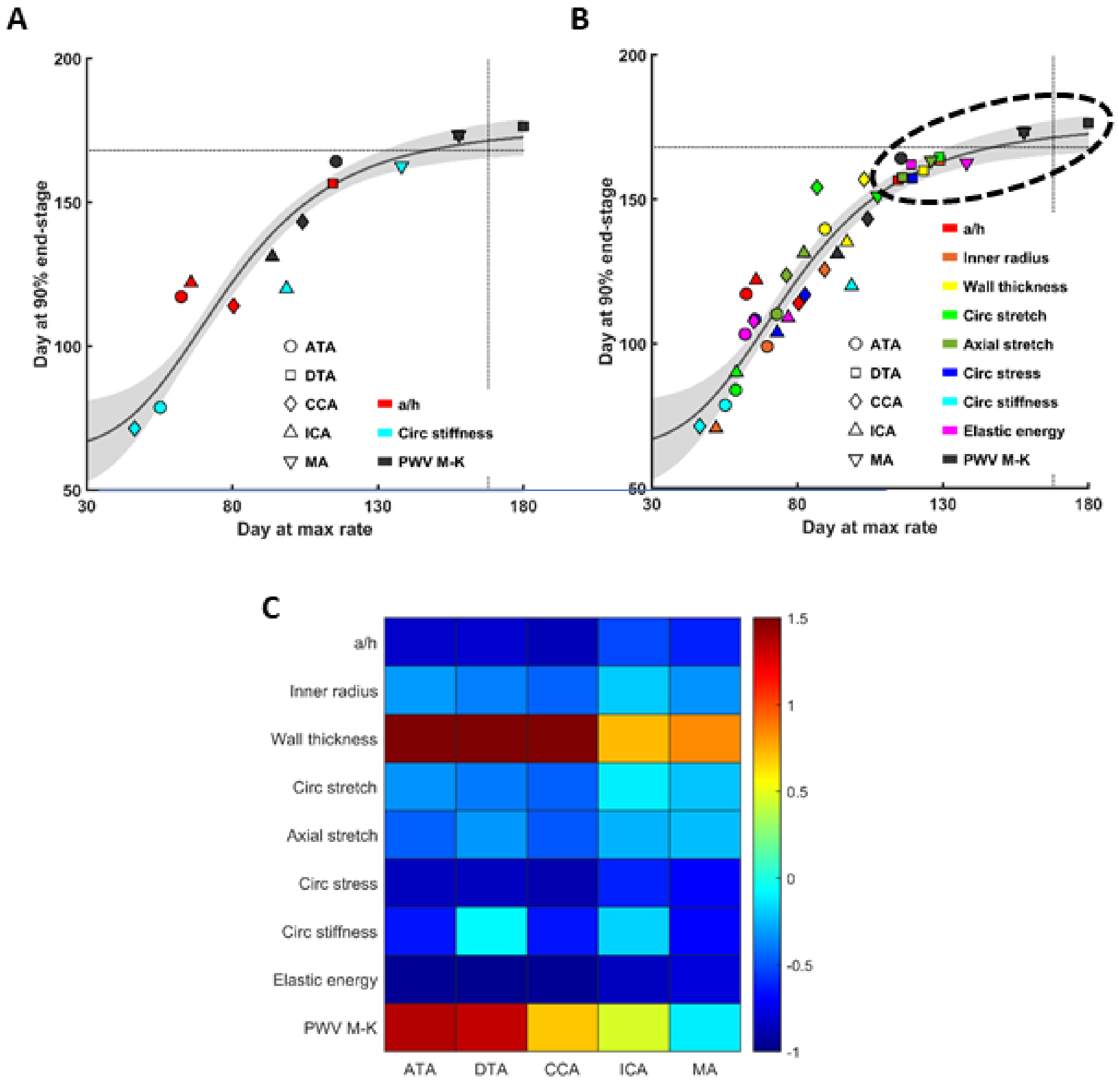


Figure S9. (A, B) Visual summary of progressive changes in five systemic arterial segments (ATA, DTA, CCA, ICA, MA - symbols) in terms of up to nine different geometric and mechanical metrics (colors) represented as the postnatal day (P42 to P168) at which a value reaches 90% of its near end of life value (ordinate) versus the day at which the rate of change of that metric was maximal (abscissa). The day at which this rate was highest emerged first for circumferential stiffness (cyan) in the ATA (circle) and CCA (diamond), around 50 days, followed by changes in inner radius (orange) around 70 days and wall thickness (yellow) around 96 days. This occurred at about the same time that PWV (black) increased in the ICA (up-triangle), CCA, and ATA. Changes occurred lastly in the DTA (square) and MA (down-triangle) for all studied metrics (dashed ellipse). Panel A highlights three particularly important metrics whereas panel B shows all nine. The ellipse highlights the most dramatic changes in the near end of life period, with P168 indicated by horizontal and vertical dashed lines, with P168 the time of 50% mean survival. (C) The same data visualized at P168 for easier assessment of increases (hot colors / red and oranges) or decreases (cold colors / blues) in all nine metrics relative to age-matched controls. Note, in particular, the calculated local pulse wave velocity (PWV M-K), which shows greater changes in the more proximal aorta and lesser changes in the more distal vessels, especially the mesenteric artery (MA).

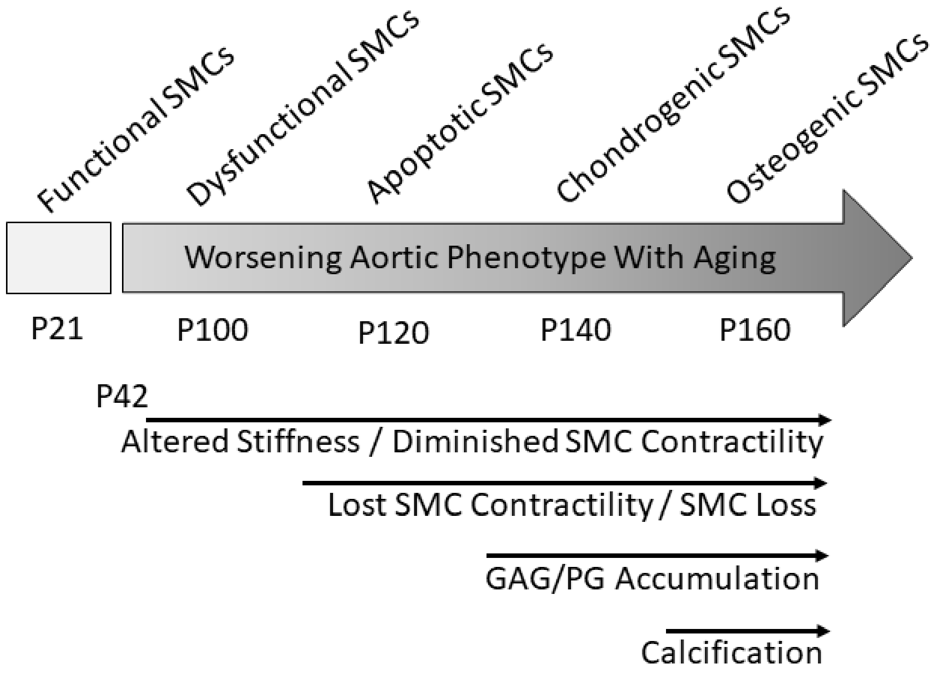


Figure S10. Schematic Summary. Smooth muscle cells (SMCs) appear to be central nodes of vascular disease progression in progeria. Schematic scale showing the worsening phenotype of the proxial aorta in progeria, with SMC phenotype indicated above and characteristics of wall structure and function below. P21, P42, P100, P120, P140, P160 are postnatal days of age, recalling that P168 was the day of mean (50%) survival. Inferences are based on the current data in combination with data previously published by Murtada et al. (2020) on SMC contractility at P100 and P140. Hence, in order, there appear to be detrimental changes in matrix turnover that affect passive properties, decreases in SMC contractility along with continued matrix turnover, a mechanical stress-mediated loss of SMCs with a subsequent phenotypic modulation by remnant SMCs leading to proteoglycan (PG) / glycosaminoglycan (GAG) accumulation and eventually calcification, all of which contribute to an increase in proximal artery stiffening that would be consistent with left ventricular diastolic dysfunction, as reported previously in humans (Prakash et al., 2018) as well as in these *Lmna*$^{G609G/G609G}$ progeria mice (Murtada et al., 2020).

### Supplemental Tables

Table S1. Geometric and passive mechanical metrics for the ascending thoracic aorta (ATA) excised from wild-type (Wt) mice at postnatal day 168 and *Lmna*$^{G609G/G609G}$ progeria (GG) mice at postnatal days 42, 100, 140, and 168, noting that the mean (50%) survival of the progeria mice is ~P168. Pressure-dependent quantities are computed at the indicated group-specific or common fixed distending pressure (P) and specimen-specific in vivo value of the axial stretch.

| | **Ascending Thoracic Aorta - ATA** | | | | |
|---|---|---|---|---|---|
| | **P42 - GG** | **P100 - GG** | **P140 - GG** | **P168 - GG** | **P168 - Wt** |
| | **n = 5** | **n = 4** | **n = 5** | **n = 5** | **n = 5** |
| **Unloaded dimensions** | | | | | |
| Wall thickness (µm) | 118 ± 3.2 | 139 ± 3.3 | 169 ± 5.9 | 180 ± 14.3 | 128 ± 4.3 |
| Outer diameter (µm) | 945 ± 13 | 1176 ± 48 | 1139 ± 19 | 1208 ± 43 | 1037 ± 30 |
| Axial length (mm) | 2.24 ± 0.14 | 2.31 ± 0.08 | 2.44 ± 0.07 | 2.58 ± 0.04 | 2.25 ± 0.10 |
| | **Systolic pressure** | | | | |
| **Loaded dimensions** | **P = 91** | **P = 101** | **P = 101** | **P = 101** | **P = 126** |
| Outer diameter (µm) | 1373 ± 34.0 | 1317 ± 46.2 | 1381 ± 16.4 | 1360 ± 18.4 | 1639 ± 38.7 |
| Wall thickness (µm) | 38 ± 1.2 | 95 ± 1.8 | 113 ± 3.7 | 143 ± 14.7 | 37 ± 1.3 |
| Inner radius (µm) | 649 ± 18.1 | 563 ± 21.8 | 577 ± 4.9 | 537 ± 6.0 | 782 ± 18.1 |
| *In vivo* axial stretch ($\lambda_z^{iv}$) | 1.95 ± 0.03 | 1.25 ± 0.02 | 1.14 ± 0.02 | 1.07 ± 0.01 | 1.96 ± 0.02 |
| *In vivo* Circumferential Stretch ($\lambda_\vartheta$) | 1.62 ± 0.03 | 1.18 ± 0.02 | 1.31 ± 0.03 | 1.19 ± 0.03 | 1.77 ± 0.03 |
| **Cauchy stresses (kPa)** | | | | | |
| Circumferential, $\sigma_\vartheta$ | 210 ± 11.6 | 80 ± 2.3 | 69 ± 1.8 | 53 ± 5.1 | 355 ± 5.0 |
| Axial, $\sigma_z$ | 319 ± 10.0 | 69 ± 3.9 | 45 ± 2.6 | 33 ± 3.0 | 327 ± 6.7 |
| **Linearized material stiffness (MPa)** | | | | | |
| Circumferential, $\mathcal{C}_{\theta\theta\theta\theta}$ | 0.91 ± 0.06 | 0.58 ± 0.05 | 0.66 ± 0.03 | 0.75 ± 0.10 | 2.76 ± 0.19 |
| Axial, $\mathcal{C}_{zzzz}$ | 1.76 ± 0.07 | 0.51 ± 0.05 | 0.49 ± 0.04 | 0.44 ± 0.02 | 0.85 ± 0.05 |
| **Stored elastic energy (kPa)** | 91 ± 2.4 | 11 ± 1.1 | 8 ± 0.5 | 4 ± 0.9 | 104 ± 1.3 |
| | **Fixed pressure** | | | | |
| **Loaded dimensions** | **P = 100** | **P = 100** | **P = 100** | **P = 100** | **P = 100** |
| Outer diameter (µm) | 1427 ± 34.3 | 1314 ± 46.2 | 1379 ± 16.5 | 1359 ± 18.5 | 1542 ± 37.8 |
| Wall thickness (µm) | 36 ± 1.1 | 95 ± 1.8 | 113 ± 3.7 | 143 ± 14.7 | 40 ± 1.3 |
| Inner radius (µm) | 678 ± 18.2 | 562 ± 21.9 | 576 ± 4.9 | 536 ± 5.9 | 731 ± 17.6 |
| *In vivo* axial stretch ($\lambda_z^{iv}$) | 1.95 ± 0.03 | 1.25 ± 0.02 | 1.14 ± 0.02 | 1.07 ± 0.01 | 1.96 ± 0.02 |
| *In vivo* circumferential stretch ($\lambda_\vartheta$) | 1.68 ± 0.03 | 1.18 ± 0.02 | 1.31 ± 0.03 | 1.19 ± 0.03 | 1.65 ± 0.03 |
| **Cauchy Stresses (kPa)** | | | | | |
| Circumferential, $\sigma_\vartheta$ | 251 ± 13.6 | 79 ± 2.3 | 68 ± 1.7 | 52 ± 5.0 | 247 ± 2.7 |
| Axial, $\sigma_z$ | 339 ± 10.5 | 69 ± 3.9 | 44 ± 2.6 | 33 ± 3.0 | 280 ± 6.9 |
| **Linearized material stiffness (MPa)** | | | | | |
| Circumferential, $\mathcal{C}_{\theta\theta\theta\theta}$ | 1.12 ± 0.07 | 0.58 ± 0.05 | 0.65 ± 0.03 | 0.73 ± 0.09 | 1.37 ± 0.06 |
| Axial, $\mathcal{C}_{zzzz}$ | 1.93 ± 0.07 | 0.51 ± 0.05 | 0.49 ± 0.03 | 0.43 ± 0.02 | 0.75 ± 0.03 |
| **Stored elastic energy (kPa)** | 101 ± 2.7 | 11 ± 1.1 | 8 ± 0.5 | 4 ± 0.9 | 85 ± 1.8 |

Table S2. Similar to Table S1 except for the descending thoracic aorta (DTA).

| | Descending Thoracic Aorta - DTA | | | | |
|---|---|---|---|---|---|
| | **P42 - GG** | **P100 - GG** | **P140 - GG** | **P168 - GG** | **P168 - Wt** |
| | **n = 5** | **n = 5** | **n = 5** | **n = 5** | **n = 5** |
| **Unloaded dimensions** | | | | | |
| Wall thickness (μm) | 97 ± 3.0 | 106 ± 3.1 | 142 ± 7.0 | 132 ± 6.1 | 98 ± 2.4 |
| Outer diameter (μm) | 703 ± 19 | 727 ± 25 | 844 ± 57 | 939 ± 21 | 796 ± 8 |
| Axial length (mm) | 4.44 ± 0.22 | 4.57 ± 0.13 | 5.78 ± 0.32 | 5.28 ± 0.08 | 4.63 ± 0.15 |
| | | | **Systolic pressure** | | |
| **Loaded dimensions** | **P = 91** | **P = 101** | **P = 101** | **P = 101** | **P = 126** |
| Outer diameter (μm) | 1025 ± 43.3 | 1033 ± 29.6 | 996 ± 27.4 | 973 ± 20.1 | 1259 ± 16.0 |
| Wall thickness (μm) | 39 ± 0.3 | 46 ± 2.2 | 99 ± 11.8 | 115 ± 7.4 | 35 ± 1.3 |
| Inner radius (μm) | 474 ± 21.5 | 470 ± 14.4 | 399 ± 13.5 | 371 ± 14.5 | 594 ± 8.2 |
| *In vivo* axial stretch ($\lambda_z^{iv}$) | 1.53 ± 0.02 | 1.44 ± 0.03 | 1.14 ± 0.02 | 1.09 ± 0.03 | 1.59 ± 0.02 |
| *In vivo* Circumferential Stretch ($\lambda_\vartheta$) | 1.62 ± 0.03 | 1.59 ± 0.04 | 1.30 ± 0.09 | 1.07 ± 0.04 | 1.75 ± 0.01 |
| **Cauchy stresses (kPa)** | | | | | |
| Circumferential, $\sigma_\vartheta$ | 147 ± 6.3 | 138 ± 7.4 | 58 ± 8.6 | 44 ± 4.3 | 286 ± 12.2 |
| Axial, $\sigma_z$ | 169 ± 3.8 | 161 ± 11.7 | 55 ± 8.8 | 67 ± 17.3 | 259 ± 11.6 |
| **Linearized material stiffness (MPa)** | | | | | |
| Circumferential, $\mathcal{C}_{\theta\theta\theta\theta}$ | 0.68 ± 0.05 | 0.71 ± 0.09 | 0.56 ± 0.11 | 1.71 ± 0.37 | 1.71 ± 0.07 |
| Axial, $\mathcal{C}_{zzzz}$ | 1.60 ± 0.04 | 1.86 ± 0.11 | 0.89 ± 0.11 | 3.69 ± 1.57 | 1.55 ± 0.14 |
| **Stored elastic energy (kPa)** | 45 ± 2.4 | 38 ± 2.2 | 9 ± 2.9 | 2 ± 0.9 | 74 ± 3.5 |
| | | | **Fixed pressure** | | |
| **Loaded dimensions** | **P = 100** | **P = 100** | **P = 100** | **P = 100** | **P = 100** |
| Outer diameter (μm) | 1060 ± 43.0 | 1029 ± 30.0 | 995 ± 27.6 | 973 ± 20.1 | 1192 ± 14.6 |
| Wall thickness (μm) | 38 ± 0.4 | 47 ± 2.2 | 99 ± 11.7 | 115 ± 7.4 | 37 ± 1.4 |
| Inner radius (μm) | 493 ± 21.3 | 468 ± 14.6 | 398 ± 13.3 | 371 ± 14.5 | 559 ± 7.6 |
| *In vivo* axial stretch ($\lambda_z^{iv}$) | 1.53 ± 0.02 | 1.44 ± 0.03 | 1.14 ± 0.02 | 1.09 ± 0.03 | 1.59 ± 0.02 |
| *In vivo* circumferential stretch ($\lambda_\vartheta$) | 1.68 ± 0.03 | 1.59 ± 0.04 | 1.30 ± 0.09 | 1.07 ± 0.04 | 1.65 ± 0.01 |
| **Cauchy Stresses (kPa)** | | | | | |
| Circumferential, $\sigma_\vartheta$ | 174 ± 6.8 | 135 ± 7.4 | 58 ± 8.4 | 44 ± 4.2 | 201 ± 9.0 |
| Axial, $\sigma_z$ | 185 ± 4.3 | 160 ± 11.4 | 54 ± 8.7 | 67 ± 17.3 | 209 ± 9.1 |
| **Linearized material stiffness (MPa)** | | | | | |
| Circumferential, $\mathcal{C}_{\theta\theta\theta\theta}$ | 0.84 ± 0.06 | 0.69 ± 0.08 | 0.55 ± 0.10 | 1.69 ± 0.37 | 1.08 ± 0.05 |
| Axial, $\mathcal{C}_{zzzz}$ | 1.91 ± 0.05 | 1.81 ± 0.10 | 0.88 ± 0.10 | 3.68 ± 1.57 | 1.16 ± 0.10 |
| **Stored elastic energy (kPa)** | 51 ± 2.4 | 37 ± 2.3 | 8 ± 2.9 | 2 ± 0.9 | 60 ± 3.2 |

Table S3. Similar to Table S1 except for the common carotid artery (CCA).

| | Common Carotid Artery - CCA | | | | |
|---|---|---|---|---|---|
| | **P42 - GG** | **P100 - GG** | **P140 - GG** | **P168 - GG** | **P168 - Wt** |
| | **n = 5** | **n = 4** | **n = 4** | **n = 5** | **n = 5** |
| **Unloaded dimensions** | | | | | |
| Wall thickness (μm) | 77 ± 1.6 | 84 ± 4.6 | 100 ± 14.7 | 102 ± 8.9 | 83 ± 2.4 |
| Outer diameter (μm) | 380 ± 11 | 412 ± 23 | 448 ± 20 | 490 ± 12 | 391 ± 14 |
| Axial length (mm) | 3.69 ± 0.19 | 3.38 ± 0.20 | 2.68 ± 0.35 | 4.39 ± 0.51 | 4.61 ± 0.13 |
| | | | **Systolic pressure** | | |
| **Loaded dimensions** | **P = 91** | **P = 101** | **P = 101** | **P = 101** | **P = 126** |
| Outer diameter (μm) | 514 ± 9.4 | 482 ± 12.7 | 472 ± 17.5 | 506 ± 24.0 | 619 ± 13.0 |
| Wall thickness (μm) | 24 ± 0.9 | 52 ± 6.9 | 79 ± 11.2 | 93 ± 9.7 | 22 ± 0.9 |
| Inner radius (μm) | 233 ± 4.0 | 188 ± 9.5 | 157 ± 15.5 | 160 ± 15.0 | 288 ± 6.3 |
| *In vivo* axial stretch ($\lambda_z^{iv}$) | 1.99 ± 0.03 | 1.24 ± 0.02 | 1.11 ± 0.02 | 1.04 ± 0.02 | 1.98 ± 0.06 |
| *In vivo* Circumferential Stretch ($\lambda_\vartheta$) | 1.62 ± 0.03 | 1.33 ± 0.11 | 1.13 ± 0.03 | 1.06 ± 0.05 | 1.96 ± 0.08 |
| **Cauchy stresses (kPa)** | | | | | |
| Circumferential, $\sigma_\vartheta$ | 118 ± 2.8 | 51 ± 8.3 | 29 ± 5.1 | 25 ± 4.1 | 226 ± 9.9 |
| Axial, $\sigma_z$ | 281 ± 17.5 | 90 ± 4.9 | 33 ± 3.7 | 215 ± 74.7 | 333 ± 36.6 |
| **Linearized material stiffness (MPa)** | | | | | |
| Circumferential, $\mathscr{C}_{\theta\theta\theta\theta}$ | 0.56 ± 0.02 | 0.45 ± 0.07 | 0.58 ± 0.14 | 0.57 ± 0.14 | 1.61 ± 0.13 |
| Axial, $\mathscr{C}_{zzzz}$ | 1.70 ± 0.05 | 1.46 ± 0.11 | 0.78 ± 0.05 | 8.98 ± 2.40 | 3.04 ± 0.38 |
| **Stored elastic energy (kPa)** | 67 ± 3.5 | 9 ± 1.7 | 2 ± 0.5 | 4 ± 2.8 | 82 ± 8.2 |
| | | | **Fixed pressure** | | |
| **Loaded dimensions** | **P = 100** | **P = 100** | **P = 100** | **P = 100** | **P = 100** |
| Outer diameter (μm) | 531 ± 9.4 | 481 ± 12.8 | 472 ± 17.5 | 506 ± 23.9 | 588 ± 14.3 |
| Wall thickness (μm) | 23 ± 0.9 | 53 ± 6.9 | 79 ± 11.3 | 93 ± 9.7 | 23 ± 0.8 |
| Inner radius (μm) | 242 ± 3.9 | 188 ± 9.5 | 157 ± 15.5 | 160 ± 14.9 | 271 ± 6.8 |
| *In vivo* axial stretch ($\lambda_z^{iv}$) | 1.99 ± 0.03 | 1.24 ± 0.02 | 1.11 ± 0.02 | 1.04 ± 0.02 | 1.98 ± 0.06 |
| *In vivo* circumferential stretch ($\lambda_\vartheta$) | 1.68 ± 0.03 | 1.32 ± 0.11 | 1.13 ± 0.03 | 1.06 ± 0.05 | 1.85 ± 0.06 |
| **Cauchy Stresses (kPa)** | | | | | |
| Circumferential, $\sigma_\vartheta$ | 140 ± 3.2 | 51 ± 8.1 | 29 ± 5.0 | 24 ± 4.1 | 159 ± 5.7 |
| Axial, $\sigma_z$ | 295 ± 17.2 | 90 ± 4.8 | 33 ± 3.7 | 214 ± 74.7 | 282 ± 30.5 |
| **Linearized material stiffness (MPa)** | | | | | |
| Circumferential, $\mathscr{C}_{\theta\theta\theta\theta}$ | 0.69 ± 0.03 | 0.44 ± 0.07 | 0.57 ± 0.14 | 0.57 ± 0.14 | 0.98 ± 0.08 |
| Axial, $\mathscr{C}_{zzzz}$ | 1.86 ± 0.05 | 1.45 ± 0.11 | 0.77 ± 0.05 | 8.98 ± 2.40 | 2.20 ± 0.27 |
| **Stored elastic energy (kPa)** | 71 ± 3.7 | 9 ± 1.7 | 2 ± 0.5 | 4 ± 2.8 | 71 ± 6.6 |

Table S4. Similar to Table S1 except for the internal carotid artery (ICA).

| | Internal Carotid Artery - ICA | | | | |
|---|---|---|---|---|---|
| | **P42 - GG** | **P100 - GG** | **P140 - GG** | **P168 - GG** | **P168 - Wt** |
| | **n = 5** | **n = 4** | **n = 4** | **n = 5** | **n = 5** |
| **Unloaded dimensions** | | | | | |
| Wall thickness (μm) | 49 ± 1.3 | 50 ± 2.4 | 54 ± 3.3 | 58 ± 2.1 | 50 ± 0.8 |
| Outer diameter (μm) | 236 ± 9 | 235 ± 5 | 289 ± 12 | 257 ± 10 | 249 ± 7 |
| Axial length (mm) | 1.12 ± 0.06 | 1.76 ± 0.15 | 1.23 ± 0.07 | 1.18 ± 0.17 | 1.37 ± 0.07 |
| | **Systolic pressure** | | | | |
| **Loaded dimensions** | **P = 55** | **P = 61** | **P = 61** | **P = 61** | **P = 76** |
| Outer diameter (μm) | 267 ± 5.7 | 258 ± 4.7 | 307 ± 2.3 | 291 ± 4.6 | 302 ± 7.4 |
| Wall thickness (μm) | 29 ± 1.2 | 36 ± 1.7 | 46 ± 3.1 | 45 ± 3.2 | 26 ± 0.6 |
| Inner radius (μm) | 105 ± 2.4 | 93 ± 2.7 | 108 ± 3.1 | 101 ± 4.5 | 125 ± 3.8 |
| *In vivo* axial stretch ($\lambda_z^{iv}$) | 1.33 ± 0.03 | 1.15 ± 0.01 | 1.07 ± 0.02 | 1.05 ± 0.02 | 1.40 ± 0.05 |
| *In vivo* Circumferential Stretch ($\lambda_\vartheta$) | 1.28 ± 0.04 | 1.20 ± 0.01 | 1.13 ± 0.08 | 1.25 ± 0.07 | 1.39 ± 0.02 |
| **Cauchy stresses (kPa)** | | | | | |
| Circumferential, $\sigma_\vartheta$ | 26 ± 1.1 | 21 ± 1.4 | 20 ± 1.9 | 19 ± 1.9 | 48 ± 2.1 |
| Axial, $\sigma_z$ | 44 ± 6.0 | 21 ± 1.1 | 23 ± 5.2 | 28 ± 8.0 | 65 ± 8.4 |
| **Linearized material stiffness (MPa)** | | | | | |
| Circumferential, $\mathcal{C}_{\theta\theta\theta\theta}$ | 0.15 ± 0.00 | 0.21 ± 0.04 | 0.41 ± 0.08 | 0.28 ± 0.03 | 0.37 ± 0.03 |
| Axial, $\mathcal{C}_{zzzz}$ | 0.38 ± 0.07 | 0.35 ± 0.10 | 0.69 ± 0.21 | 1.20 ± 0.52 | 0.57 ± 0.05 |
| **Stored elastic energy (kPa)** | 7 ± 1.1 | 2 ± 0.2 | 1 ± 0.2 | 2 ± 0.3 | 11 ± 1.2 |
| | **Fixed pressure** | | | | |
| **Loaded dimensions** | **P = 60** | **P = 60** | **P = 60** | **P = 60** | **P = 60** |
| Outer diameter (μm) | 273 ± 5.8 | 258 ± 4.8 | 307 ± 2.3 | 291 ± 4.6 | 289 ± 8.1 |
| Wall thickness (μm) | 28 ± 1.2 | 36 ± 1.7 | 46 ± 3.1 | 45 ± 3.2 | 28 ± 0.6 |
| Inner radius (μm) | 108 ± 2.5 | 93 ± 2.7 | 108 ± 3.2 | 101 ± 4.5 | 117 ± 4.3 |
| *In vivo* axial stretch ($\lambda_z^{iv}$) | 1.33 ± 0.03 | 1.15 ± 0.01 | 1.07 ± 0.02 | 1.05 ± 0.02 | 1.40 ± 0.05 |
| *In vivo* circumferential stretch ($\lambda_\vartheta$) | 1.32 ± 0.04 | 1.20 ± 0.01 | 1.13 ± 0.08 | 1.25 ± 0.07 | 1.31 ± 0.03 |
| **Cauchy Stresses (kPa)** | | | | | |
| Circumferential, $\sigma_\vartheta$ | 31 ± 1.3 | 21 ± 1.4 | 19 ± 1.8 | 18 ± 1.8 | 34 ± 1.8 |
| Axial, $\sigma_z$ | 45 ± 6.2 | 21 ± 1.1 | 23 ± 5.2 | 27 ± 8.0 | 57 ± 7.8 |
| **Linearized material stiffness (MPa)** | | | | | |
| Circumferential, $\mathcal{C}_{\theta\theta\theta\theta}$ | 0.18 ± 0.01 | 0.21 ± 0.03 | 0.40 ± 0.08 | 0.27 ± 0.03 | 0.24 ± 0.02 |
| Axial, $\mathcal{C}_{zzzz}$ | 0.39 ± 0.07 | 0.35 ± 0.10 | 0.69 ± 0.20 | 1.19 ± 0.52 | 0.45 ± 0.04 |
| **Stored elastic energy (kPa)** | 8 ± 1.2 | 2 ± 0.2 | 1 ± 0.2 | 2 ± 0.3 | 8 ± 0.9 |

Table S5. Similar to Table S1 except for the second branch of the mesenteric artery (MA).

| | Mesenteric Artery - MA | | | | |
|---|---|---|---|---|---|
| | **P42 - GG** | **P100 - GG** | **P140 - GG** | **P168 - GG** | **P168 - Wt** |
| | **n = 5** | **n = 5** | **n = 5** | **n = 5** | **n = 5** |
| **Unloaded dimensions** | | | | | |
| Wall thickness (μm) | 46 ± 2.6 | 40 ± 4.7 | 41 ± 3.1 | 55 ± 5.2 | 50 ± 1.7 |
| Outer diameter (μm) | 198 ± 10 | 199 ± 9 | 217 ± 11 | 225 ± 10 | 225 ± 8 |
| Axial length (mm) | 2.54 ± 0.18 | 3.85 ± 0.38 | 6.15 ± 1.36 | 3.44 ± 0.52 | 2.65 ± 0.28 |
| | **Systolic pressure** | | | | |
| **Loaded dimensions** | **P = 55** | **P = 61** | **P = 61** | **P = 61** | **P = 76** |
| Outer diameter (μm) | 212 ± 16.3 | 217 ± 16.9 | 232 ± 3.1 | 234 ± 8.3 | 282 ± 11.3 |
| Wall thickness (μm) | 22 ± 2.0 | 18 ± 1.2 | 27 ± 5.3 | 35 ± 4.0 | 19 ± 0.9 |
| Inner radius (μm) | 84 ± 8.1 | 90 ± 7.9 | 89 ± 4.7 | 82 ± 2.0 | 122 ± 5.4 |
| *In vivo* axial stretch ($\lambda_z^{iv}$) | 1.70 ± 0.10 | 1.73 ± 0.02 | 1.43 ± 0.14 | 1.36 ± 0.10 | 1.75 ± 0.04 |
| *In vivo* Circumferential Stretch ($\lambda_\vartheta$) | 1.25 ± 0.06 | 1.26 ± 0.09 | 1.18 ± 0.08 | 1.18 ± 0.04 | 1.51 ± 0.03 |
| **Cauchy stresses (kPa)** | | | | | |
| Circumferential, $\sigma_\vartheta$ | 28 ± 3.7 | 40 ± 3.5 | 32 ± 6.0 | 20 ± 3.0 | 65 ± 3.7 |
| Axial, $\sigma_z$ | 82 ± 29.4 | 107 ± 10.1 | 70 ± 13.3 | 33 ± 10.2 | 109 ± 8.9 |
| **Linearized material stiffness (MPa)** | | | | | |
| Circumferential, $\mathcal{C}_{\theta\theta\theta\theta}$ | 0.39 ± 0.07 | 0.22 ± 0.03 | 0.24 ± 0.04 | 0.76 ± 0.09 | 0.00 ± 0.00 |
| Axial, $\mathcal{C}_{zzzz}$ | 1.72 ± 0.43 | 1.84 ± 0.96 | 0.75 ± 0.22 | 1.85 ± 0.26 | 0.00 ± 0.00 |
| **Stored elastic energy (kPa)** | 12 ± 2.7 | 16 ± 1.2 | 9 ± 3.0 | 4 ± 1.7 | 19 ± 1.6 |
| | **Fixed pressure** | | | | |
| **Loaded dimensions** | **P = 60** | **P = 60** | **P = 60** | **P = 60** | **P = 60** |
| Outer diameter (μm) | 215 ± 16.6 | 217 ± 16.9 | 208 ± 22.6 | 234 ± 8.3 | 275 ± 11.3 |
| Wall thickness (μm) | 22 ± 2.0 | 18 ± 1.2 | 22 ± 1.9 | 35 ± 4.0 | 20 ± 0.9 |
| Inner radius (μm) | 86 ± 8.3 | 90 ± 7.9 | 82 ± 11.7 | 82 ± 1.9 | 118 ± 5.5 |
| *In vivo* axial stretch ($\lambda_z^{iv}$) | 1.70 ± 0.10 | 1.73 ± 0.02 | 1.60 ± 0.13 | 1.36 ± 0.10 | 1.75 ± 0.04 |
| *In vivo* circumferential stretch ($\lambda_\vartheta$) | 1.27 ± 0.07 | 1.26 ± 0.09 | 1.17 ± 0.08 | 1.18 ± 0.04 | 1.46 ± 0.03 |
| **Cauchy Stresses (kPa)** | | | | | |
| Circumferential, $\sigma_\vartheta$ | 32 ± 4.3 | 40 ± 3.4 | 31 ± 5.9 | 20 ± 3.0 | 48 ± 2.9 |
| Axial, $\sigma_z$ | 85 ± 29.5 | 107 ± 10.0 | 108 ± 44.9 | 33 ± 10.2 | 93 ± 8.4 |
| **Linearized material stiffness (MPa)** | | | | | |
| Circumferential, $\mathcal{C}_{\theta\theta\theta\theta}$ | 0.38 ± 0.07 | 0.22 ± 0.03 | 0.23 ± 0.04 | 0.47 ± 0.06 | 0.00 ± 0.00 |
| Axial, $\mathcal{C}_{zzzz}$ | 1.70 ± 0.42 | 1.82 ± 0.95 | 0.74 ± 0.22 | 1.33 ± 0.16 | 0.00 ± 0.00 |
| **Stored elastic energy (kPa)** | 12 ± 2.8 | 16 ± 1.2 | 14 ± 3.4 | 4 ± 1.7 | 17 ± 1.5 |

Table S6. Best-fit parameters for the relation describing the time-course of changes in multiple geometric and mechanical metrics of importance. Pulse wave velocity (PWV) was calculated using the Moens-Korteweg eqn.

| | $y = A \cdot \frac{t^D}{t^D + B^D} + C$ | | | | Max rate | 90% End-stage |
|---|---|---|---|---|---|---|
| | *A* | *B* | *C* | *D* | *t (days)* | *t (days)* |
| ***a/h*** | | | | | | |
| *ATA* | -14.5 | 70.9 | 18.1 | 4.0 | 62.4 | 117.2 |
| *DTA* | -10.51 | 121.10 | 12.2 | 6.00 | 114.5 | 156.6 |
| *CCA* | -7.9 | 83.8 | 9.8 | 7.0 | 80.4 | 114.0 |
| *ICA* | -1.4 | 74.7 | 3.7 | 4.0 | 65.8 | 122.0 |
| *MA* | -1.8 | 140.0 | 4.3 | 12.0 | 138.1 | 162.6 |
| ***Inner radius*** | | | | | | |
| *ATA* | -94.2 | 72.5 | 649.7 | 7.0 | 69.6 | 99.0 |
| *DTA* | -128.63 | 134.49 | 476.0 | 7.00 | 129.1 | 163.4 |
| *CCA* | -76.6 | 93.0 | 233.1 | 7.0 | 89.3 | 125.6 |
| *ICA* | -4.5 | 53.7 | 105.1 | 8.0 | 52.0 | 70.7 |
| *MA* | 3.3 | 53.7 | 83.7 | 8.0 | 52.0 | 70.8 |
| ***Wall thickness*** | | | | | | |
| *ATA* | 102.6 | 97.0 | 37.3 | 5.0 | 89.4 | 139.7 |
| *DTA* | 90.59 | 128.46 | 37.92 | 7.00 | 123.3 | 159.9 |
| *CCA* | 84.8 | 116.9 | 23.4 | 4.0 | 102.9 | 156.9 |
| *ICA* | 17.2 | 101.1 | 28.9 | 7.0 | 97.0 | 135.2 |
| *MA* | 21.4 | 160.0 | 21.6 | 12.0 | 157.8 | 173.4 |
| ***Circ stretch*** | | | | | | |
| *ATA* | -0.4 | 61.3 | 1.6 | 7.0 | 58.8 | 83.9 |
| *DTA* | -0.70 | 136.23 | 1.64 | 6.00 | 128.8 | 164.7 |
| *CCA* | -0.7 | 109.2 | 1.6 | 3.0 | 86.7 | 154.1 |
| *ICA* | -0.1 | 62.4 | 1.3 | 6.0 | 59.0 | 90.0 |
| *MA* | -0.1 | 113.7 | 1.3 | 6.0 | 107.5 | 151.1 |
| ***Axial stretch*** | | | | | | |
| *ATA* | -0.9 | 77.2 | 2.0 | 6.0 | 72.9 | 110.2 |
| *DTA* | -0.53 | 122.73 | 1.53 | 6.00 | 116.0 | 157.6 |
| *CCA* | -1.0 | 82.6 | 2.0 | 5.0 | 76.2 | 123.7 |
| *ICA* | -0.3 | 89.1 | 1.3 | 5.0 | 82.2 | 131.4 |
| *MA* | -0.5 | 133.1 | 1.7 | 6.0 | 125.9 | 163.4 |
| ***Circ stress*** | | | | | | |
| *ATA* | -156.5 | 70.9 | 214.9 | 5.0 | 65.4 | 108.4 |
| *DTA* | -121.74 | 124.63 | 149.13 | 7.00 | 119.6 | 157.3 |
| *CCA* | -93.5 | 86.1 | 118.2 | 7.0 | 82.6 | 116.9 |
| *ICA* | -6.7 | 76.0 | 26.5 | 7.0 | 73.0 | 103.8 |
| *MA* | -11.5 | 140.0 | 33.1 | 12.0 | 138.1 | 162.6 |
| ***Axial stress*** | | | | | | |
| *ATA* | -299.0 | 70.2 | 329.2 | 5.0 | 64.7 | 107.4 |
| *DTA* | -171.59 | 134.87 | 174.81 | 4.00 | 118.7 | 163.7 |
| *CCA* | -203.8 | 86.6 | 277.0 | 8.0 | 83.9 | 113.6 |
| *ICA* | -19.6 | 65.0 | 43.8 | 8.0 | 61.0 | 82.9 |
| *MA* | -66.5 | 160.0 | 77.2 | 12.0 | 157.8 | 173.4 |
| ***Circ stiffness*** | | | | | | |
| *ATA* | -0.3 | 57.5 | 0.9 | 7.0 | 55.2 | 78.7 |
| *DTA* | 1.54 | 160.00 | 0.63 | 12.00 | 157.8 | 173.4 |
| *CCA* | 0.0 | 49.1 | 0.6 | 6.0 | 46.4 | 71.4 |
| *ICA* | 0.2 | 100.0 | 0.2 | 12.0 | 98.6 | 120.0 |
| *MA* | -0.1 | 140.0 | 0.3 | 12.0 | 138.1 | 162.6 |
| ***Axial stiffness*** | | | | | | |
| *ATA* | -0.3 | 57.5 | 0.9 | 7.0 | 63.1 | 89.9 |
| *DTA* | -1.00 | 136.48 | 1.69 | 5.00 | 125.9 | 164.8 |
| *CCA* | 93.1 | 254.1 | 1.0 | 6.0 | 180.0 | 176.5 |
| *ICA* | 0.5 | 144.9 | 0.4 | 6.0 | 137.0 | 167.9 |
| *MA* | -10.1 | 339.1 | 1.3 | 5.0 | 180.0 | 176.1 |
| ***Elastic energy*** | | | | | | |
| *ATA* | -91.7 | 67.2 | 95.0 | 5.0 | 62.0 | 103.3 |
| *DTA* | -57.61 | 129.14 | 46.21 | 5.00 | 119.1 | 162.0 |
| *CCA* | -69.8 | 70.5 | 69.2 | 5.0 | 65.0 | 107.8 |
| *ICA* | -5.5 | 80.0 | 6.8 | 7.0 | 76.8 | 109.0 |
| *MA* | -9.6 | 140.0 | 13.4 | 12.0 | 138.1 | 162.6 |
| ***PWV M-K*** | | | | | | |
| *ATA* | 201.1 | 1078.7 | 5.0 | 2.0 | 180.0 | 170.8 |
| *DTA* | 104.43 | 241.01 | 5.15 | 6.00 | 180.0 | 176.4 |
| *CCA* | 7.7 | 108.5 | 5.3 | 7.0 | 104.1 | 143.2 |
| *ICA* | 3.9 | 97.6 | 4.6 | 7.0 | 93.7 | 131.1 |
| *MA* | 1.8 | 160.0 | 5.6 | 12.0 | 157.8 | 173.4 |
